# Improved single-cell ATAC-seq reveals chromatin dynamics of *in vitro* corticogenesis

**DOI:** 10.1101/637256

**Authors:** Ryan M. Mulqueen, Brooke A. DeRosa, Casey A. Thornton, Zeynep Sayar, Kristof A. Torkenczy, Andrew J. Fields, Kevin M. Wright, Xiaolin Nan, Ramesh Ramji, Frank J. Steemers, Brian J. O’Roak, Andrew C. Adey

## Abstract

Development is a complex process that requires the precise modulation of regulatory gene networks controlled through dynamic changes in the epigenome. Single-cell-omic technologies provide an avenue for understanding the mechanisms of these processes by capturing the progression of epigenetic cell states during the course of cellular differentiation using *in vitro* or *in vivo* models^1^. However, current single-cell epigenomic methods are limited in the information garnered per individual cell, which in turn limits their ability to measure chromatin dynamics and state shifts. Single-cell combinatorial indexing (sci-) has been applied as a strategy for identifying single-cell-omic originating libraries and removes the necessity of single-cell, single-compartment chemistry^2^. Here, we report an improved sci-assay for transposase accessible chromatin by sequencing (ATAC-seq), which utilizes the small molecule inhibitor Pitstop 2™ (scip-ATAC-seq)^3^. We demonstrate that these improvements, which theoretically could be applied to any *in situ* transposition method for single-cell library preparation, significantly increase the ability of transposase to enter the nucleus and generate highly complex single-cell libraries, without altering biological signal. We applied sci-ATAC-seq and scip-ATAC-seq to characterize the chromatin dynamics of developing forebrain-like organoids, an *in vitro* model of human corticogenesis^4^. Using these data, we characterized novel putative regulatory elements, compared the epigenome of the organoid model to human cortex data, generated a high-resolution pseudotemporal map of chromatin accessibility through differentiation, and measured epigenomic changes coinciding with a neurogenic fate decision point. Finally, we combined transcription factor motif accessibility with gene activity (GA) scores to directly observe the dynamics of complex regulatory programs that regulate neurogenesis through developmental pseudotime. Overall, scip-ATAC-seq increases information content per cell and bolsters the potential for future single-cell studies into complex developmental processes.

## Main

Recent methodical advances have enabled the preparation of thousands of single-cell-omics libraries simultaneously. Using the general sci-framework, single-cell library generating methods have been developed to measure accessible chromatin^5^, genomic sequence variation^6^, transcription^7^, chromatin folding^8^ and DNA methylation^9^. Specifically, sci-ATAC-seq enables the interrogation of open chromatin regions, which are predominantly active promoters and enhancers, and make up between 1-4% of the genome^10^. In sci-ATAC-seq, generation of sequencing library molecules is selective towards regions of open chromatin due to the steric hindrance caused by DNA-bound proteins such as histones, on the hyperactive derivative of the cut-and-paste Tn5 transposase^11^. The sci-ATAC-seq platform was recently utilized to produce whole-organism maps^2^, demonstrating the throughput and power of the technique. We reasoned that by increasing the information content generated per cell without limiting the number of cells assayed in the sci-ATAC-seq platform, we could better characterize the dynamics of complex tissue-level development.

The effectiveness of transposition, and thus sci-ATAC-seq library complexity, is limited by several factors, including: i) the amount of mitochondrial DNA, which can potentially lead to non-genomic sequence library molecules; and ii) the hindrance of cell and nuclear membranes to transposase nuclear entry. Recent adaptations to single-cell ATAC-seq protocols have worked to address both of these issues by the addition of various detergents and additional nuclear washing steps^12,13^. However, treatment with harsh detergents increases the potential of nuclear lysis, which removes the inherent compartmentalization of *in situ* transposition. Therefore, we sought to develop an alternative approach to disrupt the ultra-structure of the nuclear pore complex (NPC), while still maintaining nuclear integrity and compartmentalization.

The small molecule inhibitor *N*-[5-(4-Bromobenzylidene)-4-oxo-4,5-dihydro-1,3-thiazol-2-yl]naphthalene-1-sulfonamide, which is marketed as Pitstop 2™ (Abcam), has been demonstrated to disrupt the phenylalanine-glycine rich proteins binding within the NPC^3^. Unlike other aliphatic alcohol and detergent additions, Pitstop 2 specifically disrupts the NPC and not the nuclear membrane lipid bilayer^3^. Furthermore, washout of Pitstop 2 alleviates NPC disruption and reestablishes nuclear integrity^3^. Both of these factors made Pitstop 2 an ideal candidate small molecule for *in situ* protocols that require maintenance of nuclear integrity in a controlled and reversible manner.

We hypothesized that the addition of Pitstop 2 would increase the permeability of the NPC to macromolecules and allow for increased transposase nuclear occupancy of individual nuclei. As a consequence, a greater number of transposition events and sequencing library complexity could be achieved, relative to established methods. To measure transposase-nuclear co-localization at a population level, we performed an experiment in the well-characterized human lymphoblastoid cell line, GM12878, using varying concentrations of Pitstop 2 and transposase complexed with oligonucleotides containing a Cy5 fluorophore. Relative fluorescence of nuclei and transposase were measured by flow cytometry. We found that Pitstop 2 had an additive effect for increasing transposase-nuclear co-localization up to concentrations of 70 μM, but with the highest concentration (90 μM) showing a negative effect (Fig. 1a).

**Fig 1.**
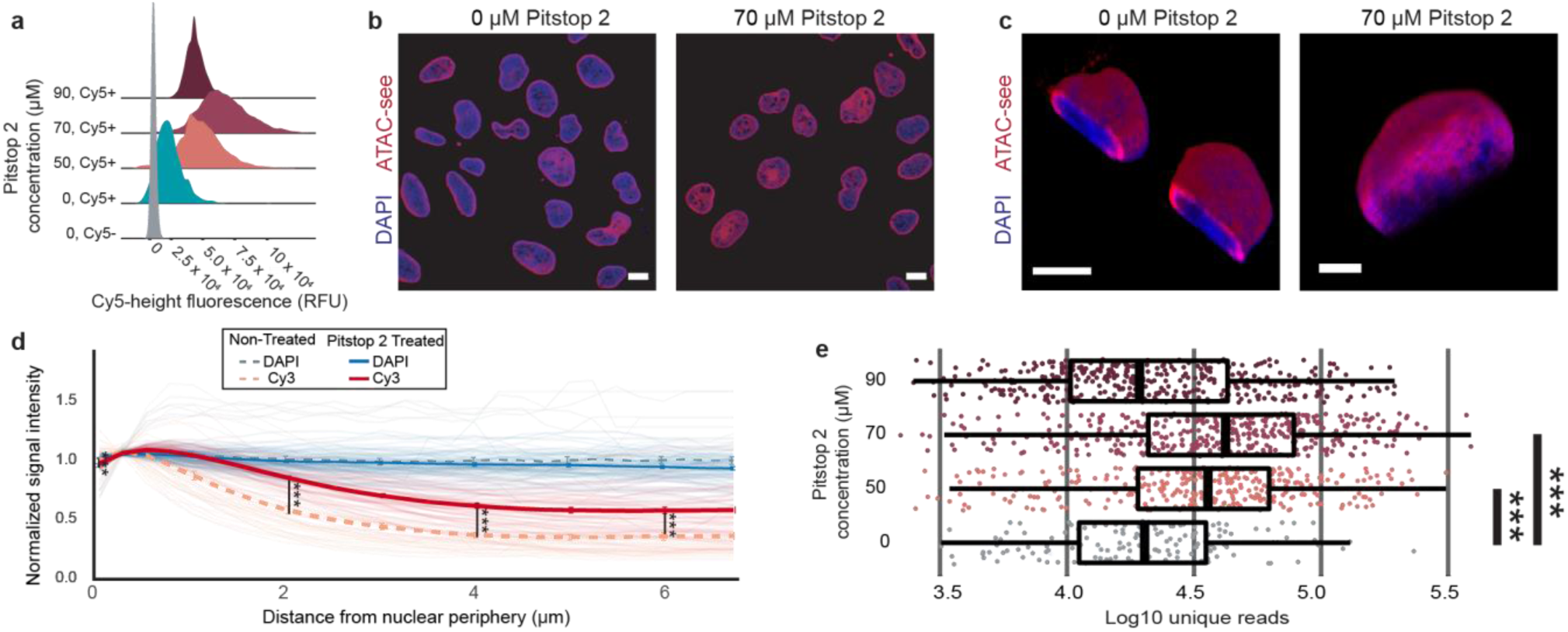
Pitstop 2 increases single-cell library complexity by increased Transposase-nuclear occupancy. **a**, Relative Cy5 fluorescence of GM12878 nuclei quantified via flow cytometry following treatment of Pitstop 2 and Cy5-fluorescently tagged transposase. Conditions contain an average of 10,526 events (minimum n=3,312 events). **b**, Representative fluorescent imaging of U2O2 nuclei showing the co-localization of Cy3-fluorescently tagged transposase and DAPI-stained nuclei. The left image shows non-Pitstop 2 treated nuclei and right side shows 70 μM Pitstop 2 treated nuclei. Scale bar represents 10 μm. **c**, Representative Z-stacked images of two nuclei without stop 2 treatment (left; scale bar is 8 μm). One representative nucleus imaged following Pitstop 2 treatment (right; scale bar is 5 μm). **d**, Quantification of Cy3 fluorescent skewness with respect to nuclear centers. Quantitative comparison of Cy3-fluorescently tagged transposase nuclear occupancy in 70 μM Pitstop 2 treated nuclei (red, n = 53) and untreated nuclei (orange, n = 42). Mean normalized signal intensity ± s.e. is plotted. **e**, Count of unique reads from sci-ATAC-seq GM12878 libraries generated with various concentrations of Pitstop 2. All reported p-values from Benjamini-Hochberg adjusted two-sided MW U test, **P < 5 × 10^−3^, ***P < 5 × 10^−6^.

To determine whether this observed increase in transposase-nuclei association was specifically due to increased transposase within the nucleus as opposed to non-specific effects, we complimented transposase adapters with a Cy3 fluorophore modified oligonucleotide to perform ATAC-see microscopy using nuclei from U2OS cells^14^ (Fig. 1 b,c; Extended Data Fig 1a,b). Notably, the signal intensity of Cy3 fluorescence was significantly higher near the nuclear periphery in non-treated samples, confirming the nuclear membrane acts as a barrier to entry (Fig. 1d, p-value ≤ 1.4 × 10^−3^, Mann-Whitney U test, Benjamini-Hochberg corrected). In contrast, Pitstop 2 treated nuclei exhibited increased Cy3 fluorescent signal localized within the interior sections of the nuclei (p-value ≤ 1.8 × 10^−9^, 4.8 × 10^−10^, and 2.3 × 10^−8^ at 2, 4, and 6 μm, respectively). At Pitstop 2 concentrations exceeding 70 μM, we observed a loss of nuclear integrity and eventual genomic leakage (Extended Data Fig. 1c,d). These data suggest addition of Pitstop 2 can dramatically increase the amount of transposase within nuclei. Moreover, a Pitstop 2 concentration of 70 μM balances the effects of increased nuclear occupancy while preserving nuclear stability.

We next addressed the practical implications of increased transposome occupancy within intact nuclei on single-cell sequencing libraries. We generated scip-ATAC-seq libraries from GM12878 cells, again using a range of Pitstop 2 concentrations. We found significant increases in unique library molecules per cell, which tracked with the additive effects noted for nuclear occupancy (Fig. 1e; p-value ≤ 3.5 × 10^−9^, 7.6 × 10^− 14^, for 50 μM and 70 μM treatment, respectively; Mann-Whitney U test, Benjamini-Hochberg corrected). We found that the addition of 70 μM Pitstop 2 to the protocol increases library complexity by an average of 2.11-fold. Notably, this effect persists if libraries are down-sampled to matched sequencing effort (Extended Data Fig. 2a; p-value ≤ 2.0 × 10^−16^, 5.2 × 10^−23^, for 50 μM and 70 μM treatment, respectively; Mann-Whitney U test, Benjamini-Hochberg corrected). We observed no adverse effects from Pitstop 2 on the distribution of library molecule size, and little decrease in the fraction of reads within peaks (FRIP, peaks defined as genomic regions of read-pileups, Extended Data Fig. 2b-c, decrease in median FRIP per cell of 14.4% for 70 μM treatment). At the library complexity we are achieving, we obtain roughly 7.11 ± 4.52% (mean ± s.d.) of the possible sites per cell as defined by all sites contained within our full GM12878 read set (Extended Data Fig. 2d; Supplementary Table 1). This suggests that the use of Pitstop 2 is successful in increasing the unique library molecules per cell while not disrupting the epigenomic configuration.

With the improved read counts per cell provided by scip-ATAC-seq, we sought to characterize a complex sample with actively forming cell types through differentiation. Brain organoids are a powerful model system to study human neurodevelopment *in vitro*^4^. Data from bulk and single-cell RNA-seq, H3K27ac ChIP-seq (chromatin-immunoprecipitation and sequencing), and bulk DNA methylation analysis, demonstrate that these models are strongly correlated with similar data from primary human fetal brain samples ranging from the early to midfetal period (post-conception weeks 9-24)^4,15,16^. Specifically, forebrain-like organoids derived from induced pluripotent stem cells (iPSCs) mimic the early stages of human corticogenesis and lamination, wherein proliferating radial glia cells in the ventricular zone generate a pool of progenitors^4,17,18^. From these radial glia cells, tightly regulated transcription factors drive either continued proliferation, or neurogenesis, when the radial glia or its intermediate progenitor differentiates into neurons^19^. However, our understanding of the temporal dynamics of non-coding regions and regulatory sites within this critical timeframe is lacking. For these reasons, we chose a forebrain-like organoid model system for leveraging our scip-ATAC-seq method on studying epigenomic dynamics^4^.

We differentiated forebrain-like organoids from human iPSCs for up to 90 days *in vitro* (DIV) using a previously described miniature bioreactor protocol with modifications to increase organoid uniformity (Online Methods, Fig. 2a, Supplementary Tables 2-4)^4^. Subsets of organoids were collected from an initial pool of 96 and characterized by their expression of cortical markers at multiple time points. Similar to previous results^4^, these forebrain-like organoids mimicked the *in vivo* developmental processes by developing multi-layered structures resembling the ventricular zone comprised of SOX2+/PAX6+ progenitors, subventricular zone comprised of EOMES+ (aka TBR2) intermediate progenitors and cortical plate, where we observe almost exclusive expression of layer-specific neuronal markers such as TBR1, CTIP2 (aka BCL11B), and SATB2 (Fig. 2a)^4^. Additionally, formation of post-mitotic neurons followed the expected step-wise temporal order of layer-specific neurogenesis, indicated by generation of TBR1+ layer 6 neurons prior to layer 5 CTIP2+ neurons, followed by SATB2+ and CUX1+ upper layer neurons (Extended Data Figs. 3 and 4).

**Fig 2.**
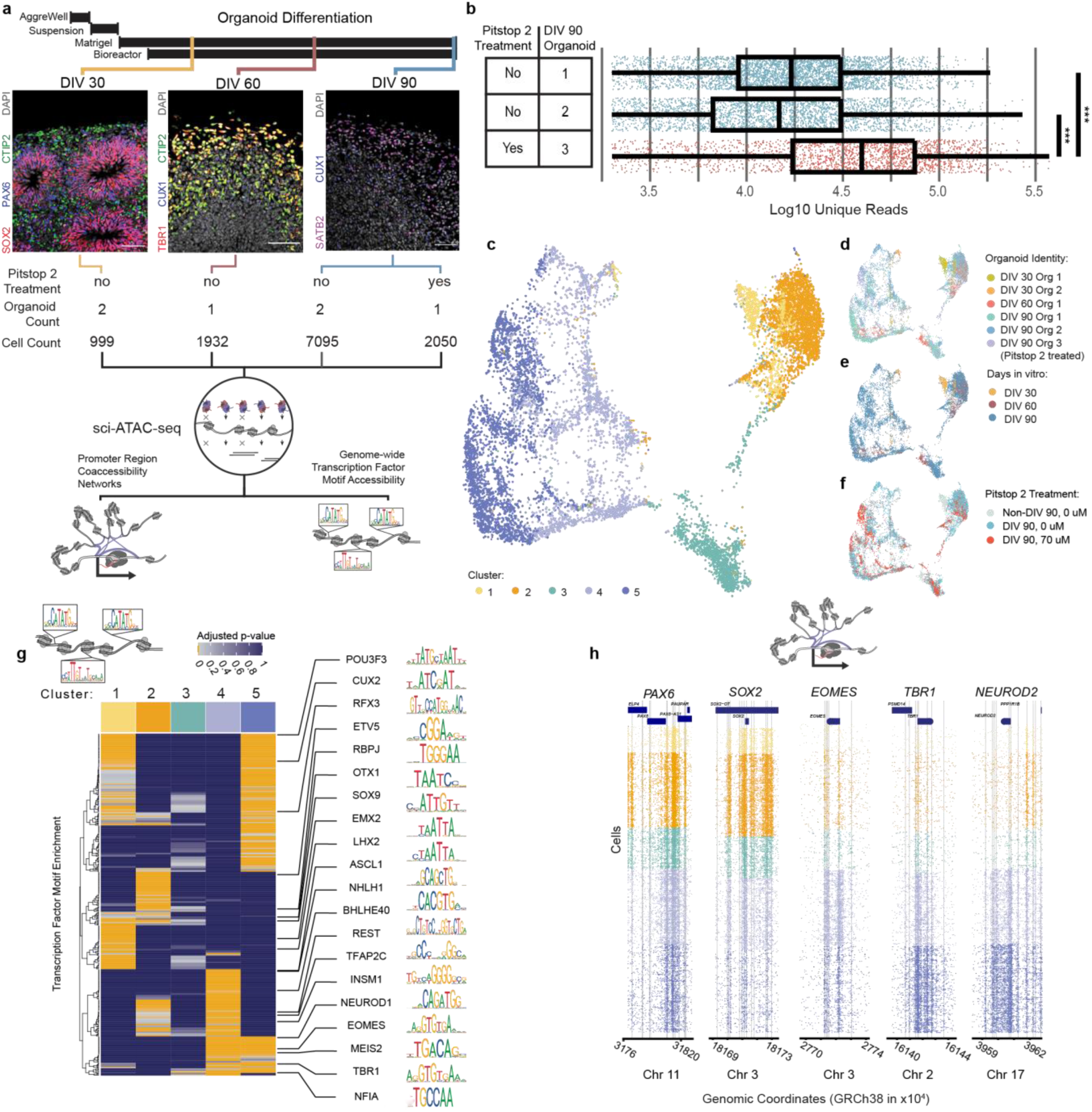
Cortical organoid cell type identification. **a**, Differentiation protocol timeline for organoid generation (top). Example tmmunohistochemistry showing positive markers for corticogenic cell type likeness. Scale bar is 50μm (middle, see Extended Data Figs 2,3). Sampled organoids and summary statistics for sci-ATAC-seq and scip-ATAC-seq protocol (bottom). **b**, Improvement of library complexity by treatment of 70μM Pitstop 2 treatment. DIV 90 organoid sourced single-cell comparison for non-treated and treated organoids prior to library generation. All reported p-values from Benjamini-Hochberg adjusted two-sided MW U test, ***P < 5×10^−6^. **c**, UMAP projection of cisTopic topics, single libraries are colored by phenogram defined clusters. **d-f**, UMAP projection colored by organoid source, DIV, and Pitstop 2 treatment, respectively. **g**, Enrichment of transcription factor motifs across sites differentially accessible across clusters. Motif enrichments were determined through hypergeometric tests following motif scanning of differentially accessible peaks per cluster. Heatmap is hierarchically clustered by correlations in Benjamini-Hochberg corrected p-values. Selected transcription factors are accompanied by their JASPAR motif. **h,** Read plots of genomic locations. Each row is a single library and all points reflect a uniquely mapped read. Vertical gray lines denote accessibility peaks. Points are colored by clusters to be consistent with panel **c**.

We performed sci-ATAC-seq and scip-ATAC-seq to generate 12,076 quality control (QC)-passing single-cell ATAC-seq profiles from two DIV 30 organoids, one DIV 60 organoid, and three DIV 90 organoids (Fig. 2a). We found that Pitstop 2 treatment significantly increased the library complexity with an average improvement of 2.6-fold, relative to DIV 90-matched organoids (Fig. 2b; p-value ≤ 3.5 × 10^−176^; Mann-Whitney U test). This increased complexity persisted when controlling for uneven sequencing effort (Extended Data Fig. 5a; p-value ≤ 4.8 × 10^−198^; Mann-Whitney U test). We further recapitulated our results showing little to no epigenome-wide disruption due to Pitstop 2 treatment (Extended Data Fig. 5 b-c, median FRIP decrease of 10.2%). Cells sourced from the Pitstop 2 treated organoid had relatively high genomic coverage, with a median of 6.9% of possible reads covered from the full read set (Extended Data Fig. 5d; Supplementary Table 5).

Using data from all cells passing QC, we identified accessible regions (peaks) across the genome. Notably, of the 125,645 called peaks, 68.5% (86,061) fell within previously defined enhancer or promoter regions^20^. Further, 46.6% (58,511) of peaks fell within previously described human adult cortex or organoid active enhancer regions, defined orthogonally with histone 3 lysine residue 27 acetylation (H3K27ac) enriched peaks (Supplementary Table 6)^13,16^. Of the remaining novel, non-intersecting peaks (58,511), 2.3% (1,351) fell in promoter regions and the remaining 97.7% (57,160) were uncharacterized distal elements. To assess the biological validity of our novel peak set, we generated pairwise correlation values for the co-occurrence of transcription factor binding motifs in our known and novel peak sets. The transcription co-occurrence matrixes between known and novel peaks were themselves highly correlated (Pearson’s 0.98), suggesting that novel distal elements recapitulate genomic motif groupings of known regulatory elements (Extended Data Fig. 6).

We used the full set of peaks and performed dimensionality reduction through the use of cisTopic^22^, a machine learning approach which defined an optimal 67 “topics” based on shared peak accessibility, producing a matrix of cells by topic weights (Extended Data Fig. 7). This matrix was then used to identify five clusters using the Louvain-based *PhenoGraph* tool^23^. We next projected the clusters into 3-dimensional space for visualization using uniform manifold approximation positioning^24^ (umap, Supplementary Table 5, interactive 3D plot: Supplementary File 1). We noted cluster biases dependent on organoid differentiation state but Pitstop 2 treatment did not interfere with DIV 90 organoid clustering position (Fig. 2c-f).

We next sought to characterize the cell type and differentiation state of the five clusters by identifying differentially accessible (DA) peaks unique to each, and detecting transcription factor motif enrichment within these cluster-specific peaks^25^. This enabled us to assign broad cell type identities to clusters 1 through 5 based on their progression through corticogenesis according to their respective sets of enriched motifs (Supplementary Table 7)^19^. For example, cluster 1 and 2 peaks showed enrichment in motifs for ETV5, a proliferating radial-glia associated transcription factor, and motifs for OTX1, which marks a neuroepithelium to forebrain commitment. Clusters 4 and 5 showed motifs associated with the NEUROD and T-box family, such as NEUROD2, EOMES (aka TBR2) and TBR1, suggesting they were populated by post-mitotic neurons (Fig. 2g)^4,26,27^. Cluster 3 had few unique DA peaks, suggesting it was a transition state between clusters 1 and 2, and clusters 4 and 5. To test this hypothesis, we performed pairwise cluster DA peak calling and motif enrichment. We found that cluster 3 is enriched in proliferative/forebrain specification marker motifs relative to clusters 4 and 5 (e.g., LHX2 and EMX2) and enriched in neurogenic marker motifs relative to clusters 1 and 2 (e.g., NFIA and TFAP2C; Extended Data Fig. 8). Cluster 3 showed specific motif enrichments for STAT3, POU2F1 and RELA, supporting the notion that it contained later born outer radial-glia. Emergence of HOPX+ outer radial glia marked cells was previously reported at latter time points (e.g., DIV 56)^4^. We found a paucity of DIV 30 sourced cells in cluster 3, further supporting the emergence of cluster 3 in later stages of cortiogenesis (Fig. 2e).

We further examined the accessibility levels at proximal elements for a set of genes with a known window of activity in corticogenesis based on the assumption that *cis*-acting elements should have increased accessibility when a gene is active^13,28^. We examined accessibility levels at proximal elements for a set of genes with a known window of activity in corticogenesis. Clusters 1, 2 and 3 showed a higher density of proximal reads for neural progenitor genes such as SOX2 or PAX6^26,27,29^. Cluster 4 showed the highest proximal read density around the EOMES region, suggesting it contains cells in an intermediate progenitor like epigenomic state. Both cluster 4 and 5 showed increased read density around deep layer neuronal cortex markers such as TBR1 and NEUROD2^4,26,27^, indicating that these marks are activating as cells are exiting the basal-progenitor like state, with an overlap of the two marker sets during the transition (Fig. 2h). These assessments further confirmed our progressive cluster assignments through corticogenesis. It is notable that due to similar or shared DNA binding motifs of many classes of transcription factors, the combination of promoter region accessibility and genome-wide transcription factor motif presentation jointly inform interpretation. For instance, cluster 1, 2, and 5 showed PAX6 DNA binding motif enrichment (Fig. 2h); however, the promoter region accessibility reveals the highest potential transcription in clusters 1 and 2.

Reasoning that all cells within the organoid samples were sourced from a shared stem-like origin and that clustering recapitulates the stereotyped patterns of corticogenesis, we used *monocle3* to generate a pseudotemporal ordering of cells – a trajectory of epigenomic changes during differentiation (Fig. 3a)^30^. We rooted the trajectory within cluster 1, populated almost solely by DIV 30 organoid-sourced cells most similar to the primodial neuroepithelia (Fig. 2e). We then used *chromVAR* to analyze the putative progression of transcription factor activities, as measured by the accessibility of the associated DNA motif genome-wide^31^. We found that multiple waves of transcription factor motif opening occur during organoid differentiation, including known patterns that have been observed in single-cell transcriptomic studies of murine corticogenesis, and bulk ATAC-seq in fetal human samples^32,33^. The earliest waves can be explained by a neuroepithelial-like state, showing relative increases in transcription factor motifs associated with telencephalic commitment or symmetric division in proliferating radial glia cells, such as OTX2 or EMX2^34^. Progressive waves follow known programs of transcription factors linked to corticogenesis, with many transcription factor waves spanning cluster boundaries (Fig. 3b). Transient increases in transcription factors associated with radial-glial proliferation, e.g., POU3F3 (aka BRN1), EOMES and NFIX, precede the waves of transcription factors such as MEF2C, NEUROD6, NEUROG2, and BACH2, which are linked to neuronal migration and maturation^32,33^.

**Fig 3.**
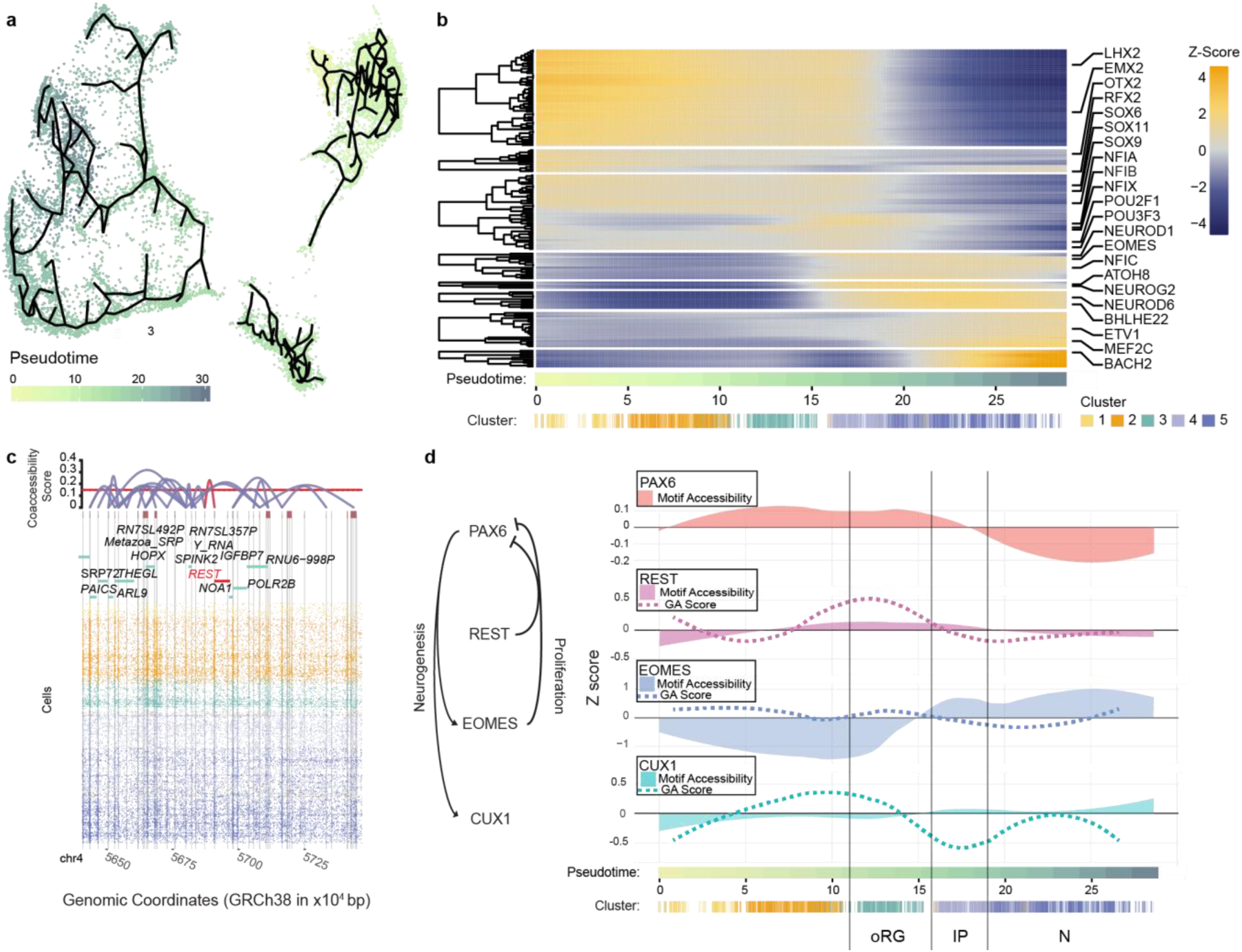
Chromatin dynamics of cortical organoid differentiation. **a**, Projection by umap of cells overlaid with minimal spanning tree generated via *monocle*^30^ and used to calculate pseudotime through differentiation. Cells are colored by pseudotime. **b**, Transcription factor motif opening through pseudotime. Motif accessibility is calculated cell-wise, ordered through pseudotime, and then smoothed by LOESS function. Rows are clustered based on Spearman’s correlation. Select transcription factor motifs are shown to the right. **c**, Co-accessibility of peaks surrounding the gene region for chromatin remodeler REST. (top) co-accessibility linkages between aggregated peaks (shown as red blocks) with height of linkage signifying co-accessibility score. (bottom) Read plot of unique reads per cell plotted in the aligned genomic region. Vertical gray bars are *de novo* called peaks. Points are unique reads aligned to the genome. Points are colored to match cluster identity as in panel b. **d**, LOESS smoothed transcription factor motif accessibility (filled curve) and gene activity score (stipled curve) through pseudotime for four factors characterized in neurogenesis. Pseudotime is demarcated for visualization by hypothesized cell types, outer radial glia (oRG), intermediate progenitors (IP) and neurons (N). GA score for PAX6 could not be generated since peak anchor region could not be unambiguously assigned (See Online Methods).

We observed a splitting of neuronal trajectories through cluster 4 and 5, suggestive of a persisting intermediate-like epigenome before leading to pathways of maturation (Fig. 3a). Tracing the accessibility of peaks defined within cisTopic-defined topics (2732+/-289 peaks per topic), we rebuilt the changes in topic accessibility as differentiation forked (Extended Data Fig. 9). Cells nearest the split were defined by topics with peaks enriched for marker motifs of intermediate progenitors, such as NHLH1, REST and ASCL1. From this branch point, three major lineages formed with the lower and upper branches showing the formation of newborn excitatory neurons with marker enrichments such as MEF2C and NEUROD1. The middle branch showed a persistence of an epigenome similar to intermediate progenitor markers with the addition of inhibitory neuron marker motifs such as MEIS1^19^.

Single-cell ATAC-seq data provides a unique opportunity to identify tightly coordinated co-accessible regulatory elements using a recently-described strategy, *cicero*^35^. We applied this analysis to our dataset, which enabled us to interconnect 98,947 of our 125,645 peaks (78.8%). We generated 149,010 correlations with coaccessbility scores ≥ 0.15, a threshold consistent with previous literature^2,13,35^. Of these, 0.5% (706) were promoter-promoter links, 7.6% (11,289) distal-promoter links, and 92% (137,015) distal-distal links. We next identified 3,850 cis-coaccessibility networks (CCANs), which incorporated 53.9% (67,688) of our total peaks. These CCANs included distal-promoter interactions for a number of key genes during corticogenesis, including *REST, LHX2, NFIB*, and *NEUROG2* (Fig. 3c, Extended Data Fig. 10a). Taken one step further, *cicero* offers a framework to utilize chromatin accessibility signal at peaks linked to promoter regions to approximate a gene activity score (GA score). We produced GA scores on local cell aggregates to improve signal at any given gene. Notably, we observed increased GA scores for genes corresponding to their expected progression through cortical development. This includes genes for transcription factors we were able to assess previously such as *SOX2* in cluster 1 and 2, and *NEUROG2* in clusters 4 and 5. We were also able to see the GA score progression of classic corticogenic marker genes that lack DNA binding activity such as TUBB2B and NRXN1 (Extended Data Fig. 10b).

Finally, we combined GA scores of transcription factors with the accessibility scores of their respective DNA binding motifs through pseudotime to demonstrate the utility of these data for understanding the dynamics of complex regulatory programs, such as the regulation of PAX6 associated neurogenesis. PAX6 drives neurogenesis by opening up promoters and distal elements associated with neuron maturation, thus depleting the pool of progenitors^36^. REST has a unique DNA binding motif and forms a complex with PAX6 to promote continued radial glia proliferation (including outer radial glia), and inhibit PAX6 driven neurogenesis^37^. EOMES also serves to inhibit the neurogenic drive of PAX6 in intermediate progenitors that will eventually differentiate into post-mitotic superficial layer neurons, expressing factors such as CUX1^38^ (Fig. 3d).

In our combined motif accessibility and GA score data, we directly observed the dynamics of these interactions in pseudotime beginning in cluster 3, which is associated with outer radial glia (Fig. 3d, Extended Data Fig. 8). Within cluster 3, elevated PAX6 motif accessibility was commensurate with increased REST motif accessibility and GA score, consistent with the role of REST in sequestering PAX6 and silencing its neurogenic activity. As cells transition into cluster 4, we observed an increase in EOMES motif accessibility and GA scores, indicating the formation intermediate progenitor states and marks of PAX6 and REST activity decline. With the loss of REST suppression, activity marks for EOMES and CUX1 increased as neurogenesis begins predominating pseudotime. We also observed that the EOMES GA score declines during neurogenesis, while its motif accessibility remains high. This is likely due to EOMES and TBR1 sharing a common DNA binding motif, making them difficult to deconvolve^39^. We speculate that the second wave of increased EOMES motif accessibility is likely driven by TBR1, which is consistent with the increased number of TBR1+ neurons within the later DIV organoids (Extended Data Figs. 3,4).

In conclusion, we have described a single-step modification to an existing protocol and demonstrate that this simple step increases the information gained by over two-fold. We also note that this step can be incorporated to any workflow that includes *in situ* transposition of DNA, including sci-DNA-seq^6^, sci-MET^9^, and any droplet-based single-cell ATAC-seq workflow. We utilized our improved assay to characterize a burgeoning model of neurodevelopment and provide, to our knowledge, the first single-cell chromatin accessibility profile of this model. Our findings not only recapitulate known waves of epigenomic reprogramming, but produce maps of regulatory element usage at unprecedented resolution, revealing precise cascades of co-accessible patterns that incorporate novel loci.

## Author contributions

A.C.A. and F.J.S supervised all aspects of Pitstop 2 assessment on in situ tagmentation. F.J.S. and R.R. initiated Pitstop 2 experiments. R.M.M. led Pitstop 2 assessment with C.A.T. ATAC-see experiments were performed by Z.S. and X.N. A.J.F. aided in scip-ATAC-seq protocol development. B.A.D. performed all organoid differentiation, culture maintenance, and immunostaining. Organoid single-cell ATAC-seq experiments were performed by R.M.M., who led the analysis with input from K.A.T and C.A.T. Biological interpretation was performed by R.M.M., B.A.D, K.M.W., B.J.O., and A.C.A. The manuscript was written by R.M.M., B.A.D., B.J.O., and A.C.A. with input from all authors.

### Acknowledgements

We thank members of the Adey and O’Roak lab groups for their input, particularly Sara Evans, and Lolli Inman. We thank Soo-Kyung Lee for sharing reagents. We thank Luiz E. Bertassoni and Anthony Tahayeri in the OHSU School of Dentistry for help with 3D printing the Spin-O bioreactor. We thank Xuyu Qian for input into the organoid differentiation protocol. This work was supported by NIH grants (5R35GM124704-02 and 1R01DA047237-01 to A.C.A., and 5R01MH113926-02 to B.J.O).

## Data and software availability

A wrapper for the entire analysis (scitools) as well as custom scripts used will be made available on github.com/adeylab/organoid. All sequence read data for CIRM lines will be made available on dbGAP, with processed data available on GEO.

---

**Extended Data Fig 1.**
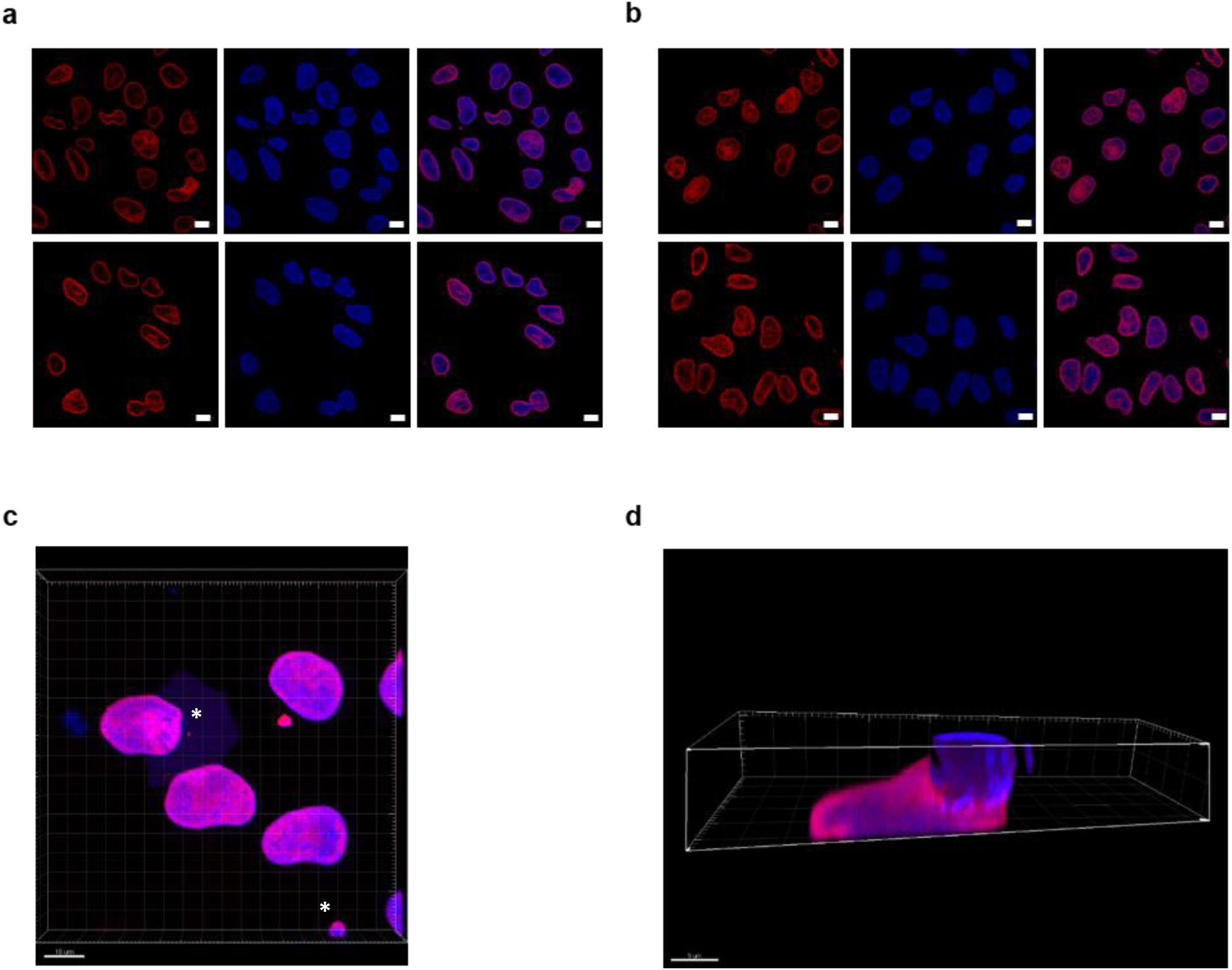
Fluorescent microscopy analysis of transposome complex. **a**, Representative fluorescent images of DAPI stained fixed nuclei with Cy3 complexed transposomes without Pitstop 2 treatment. Scale bar is 10 μm. **b**, Representative fluorescent images of DAPI stained fixed nuclei with Cy3 complexed transposomes and treated with 70 μM Pitstop 2. Scale bar is 10 μm. **c**, 3D reconstruction of DAPI and ATAC-see signal showing blend projection function of Imaris Software. Nuclear blebbing as well as bursting of a 150 μM Pitstop 2 treated nuclei and diffusion of DAPI+ DNA into the environment is visualized in 3 dimensions Scale bar is 10 μm. **d,** Cross-section of DAPI and ATAC-see signal in 150 μM Pitstop 2 treated and ruptured nuclei. Cross section created using the clipping plane function of Imaris software (Bitplane, UK) show rupture of the nucleus and spillage of DNA. Presence of ATAC-signal within the nucleus is also notable. Scale bar: 5 μm.

**Extended Data Fig 2.**
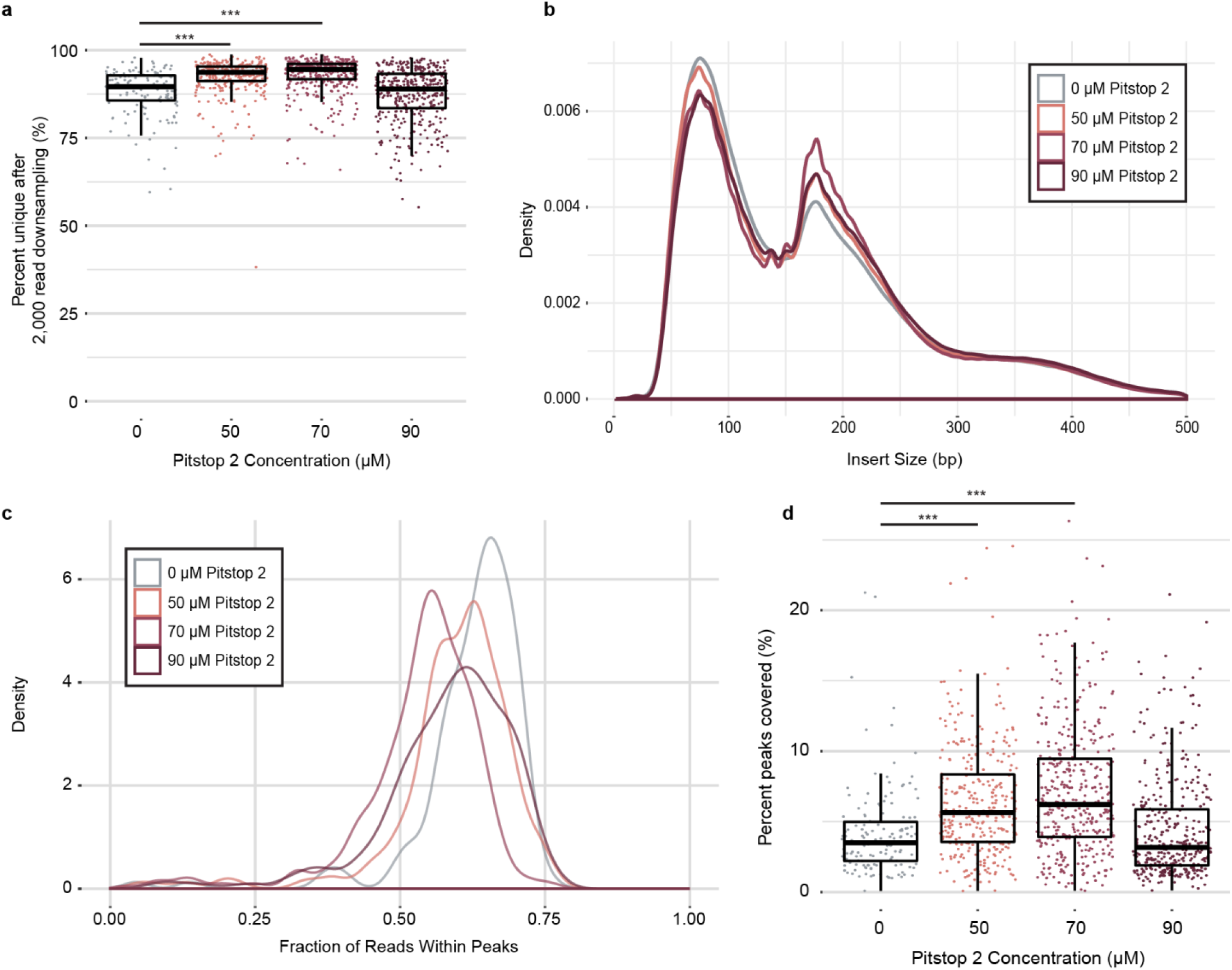
Pitstop 2 treatment increases single-cell library complexity within GM12878 cell line samples without disrupting the epigenome. **a**, Box plot of single-cell library complexities after down-sampling. Single-cell libraries were randomly sampled for 2,000 reads. The percent of unique reads per library is plotted. Significance was calculated by Pitstop 2 treatment condition to the 0 μM Pitstop 2 condition with a two-sided Mann-Whitney test. **b**, Density plot of sequencing molecule insert sizes by treatment condition. **c**, Density plot of fraction of reads overlapping with *de novo* called peaks per condition. **d**, Estimated peak coverage per cell based on read count within *de novo* called peaks. Significance was calculated by Pitstop 2 treatment condition to the 0 μM Pitstop 2 condition with a two-sided Mann-Whitney test. *** adjusted p-value ≤ 10^−4^

**Extended Data Fig 3.**
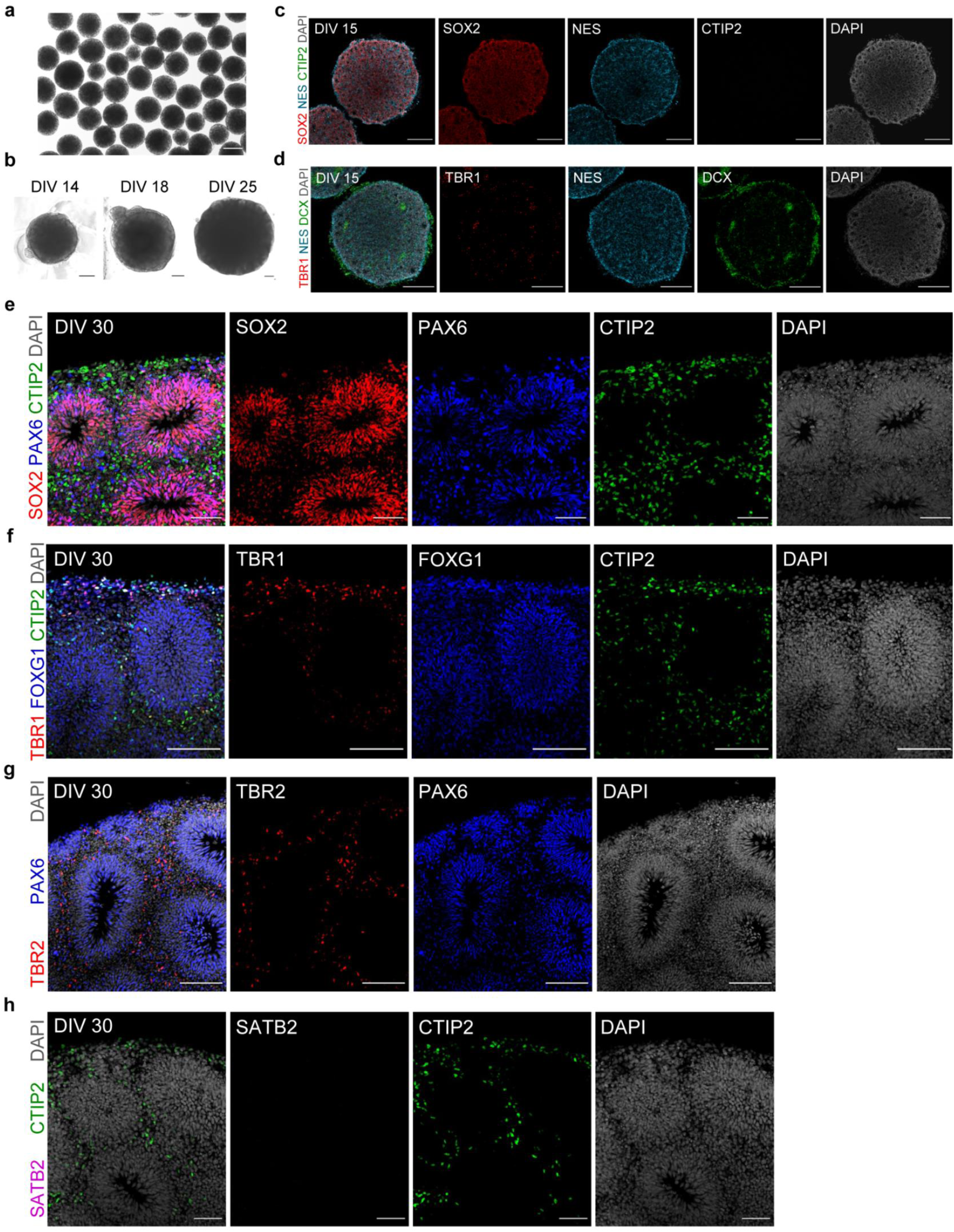
Characterization of earlier stage forebrain-like organoids. **a**, Brightfield image of days *in vitro* (DIV) 7 organoids showing uniformity in size and shape. Scale bar: 200 μm. **b,** Brightfield images of organoids at DIV 14, 18, and 25 showing growth over time. Note the increased number of neuroepithelial buds around the perimeter of DIV 25 organoids. Scale bars: 200 μm. **c-d**, Immunohistochemical characterization of organoids at DIV 15. Scale bars: 200 μm. **c**, At DIV 15, the majority of cells stain positive for the progenitor markers SOX2 and Nestin (NES) and do not yet express the layer 5 marker CTIP2. Scale bars: 200 μm. **d**, DIV 15 organoids stained for NES, DCX, and TBR1. Scale bars: 200 μm. At DIV 15, organoids are mainly comprised of SOX2+/NES+ progenitors (**c**) with patches of newly born DCX+/TBR1+ layer 6 neurons (**d**), which are the first to be born during cortical neurogenesis. **e-h**, Expanded immunohistochemical characterization of organoids at DIV 30. **e**, DIV 30 organoid with image panels show individual staining of SOX2, PAX6, and CTIP2. Scale bars: 50 μm. As seen in Fig 2a. **f**, DIV 30 organoid immunostained for the deep layer neuron markers TBR1 and CTIP2, in addition to FOXG1, a general marker of forebrain development. The subsequent expression of CTIP2 (layer 5) after TBR1 (layer 6) mimics the stepwise order of deep layer neurogenesis *in vivo*. Scale bars: 100 μm. **g**, DIV 30 organoid immunostained for EOMES and PAX6, markers of progenitors in the subventricular zone and ventricular zone, respectfully. Scale bars: 100 μm. **h**, DIV 30 organoid immunostained for SATB2 and CTIP2. Note that at DIV 30, organoids do not yet express SATB2, a marker of upper layer neurons. Scale bars: 50 μm.

**Extended Data Fig 4.**
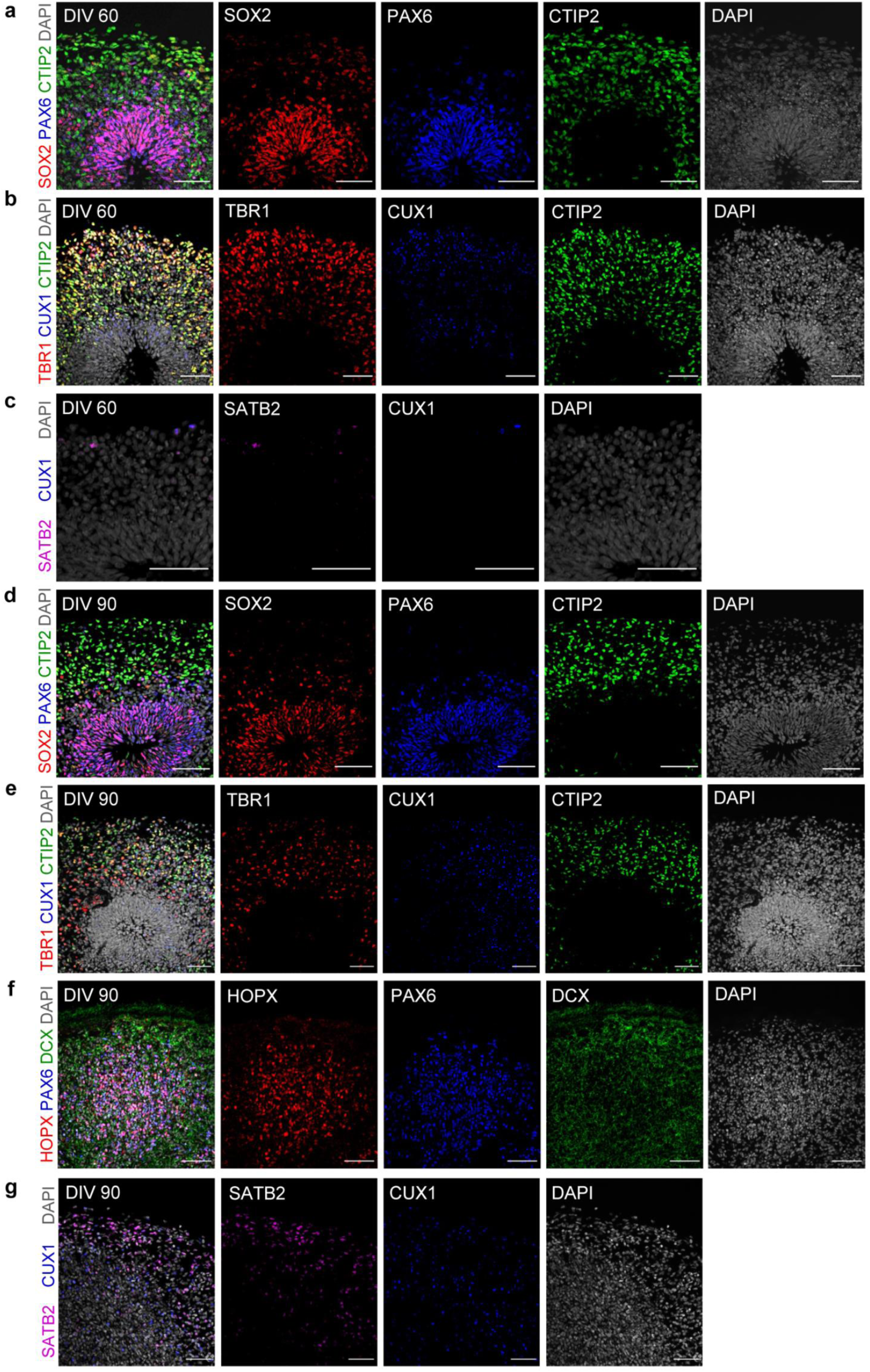
Characterization of later stage forebrain-like organoids. **a-c**, Expanded immunohistochemical characterization of organoids at days *in vitro* (DIV) 60 **a**, DIV 60 organoid immunostained for SOX2, PAX6, and CTIP2. **b**, Expanded immunohistochemical characterization of a DIV 60 organoid as seen in Fig. 2a. Image panels show individual staining of TBR1, CTIP2, and CUX1. **c**, DIV 60 organoid immunostained for CUX1 and SATB2**. d-g**, Expanded immunohistochemical characterization of organoids at DIV 90. **d,** DIV 90 organoid immunostained for SOX2, PAX6, and CTIP2. **e**, DIV 90 organoid immunostained for TBR1, CUX1, and CTIP2**. f**, DIV 90 organoid immunostained for PAX6, DCX, and HOPX, a marker outer radial glia. **g**, DIV 90 organoid as shown in Fig. 2a. Image panels show individual staining of SATB2 and CUX. Scale bars: 50 μm.

**Extended Data Fig 5.**
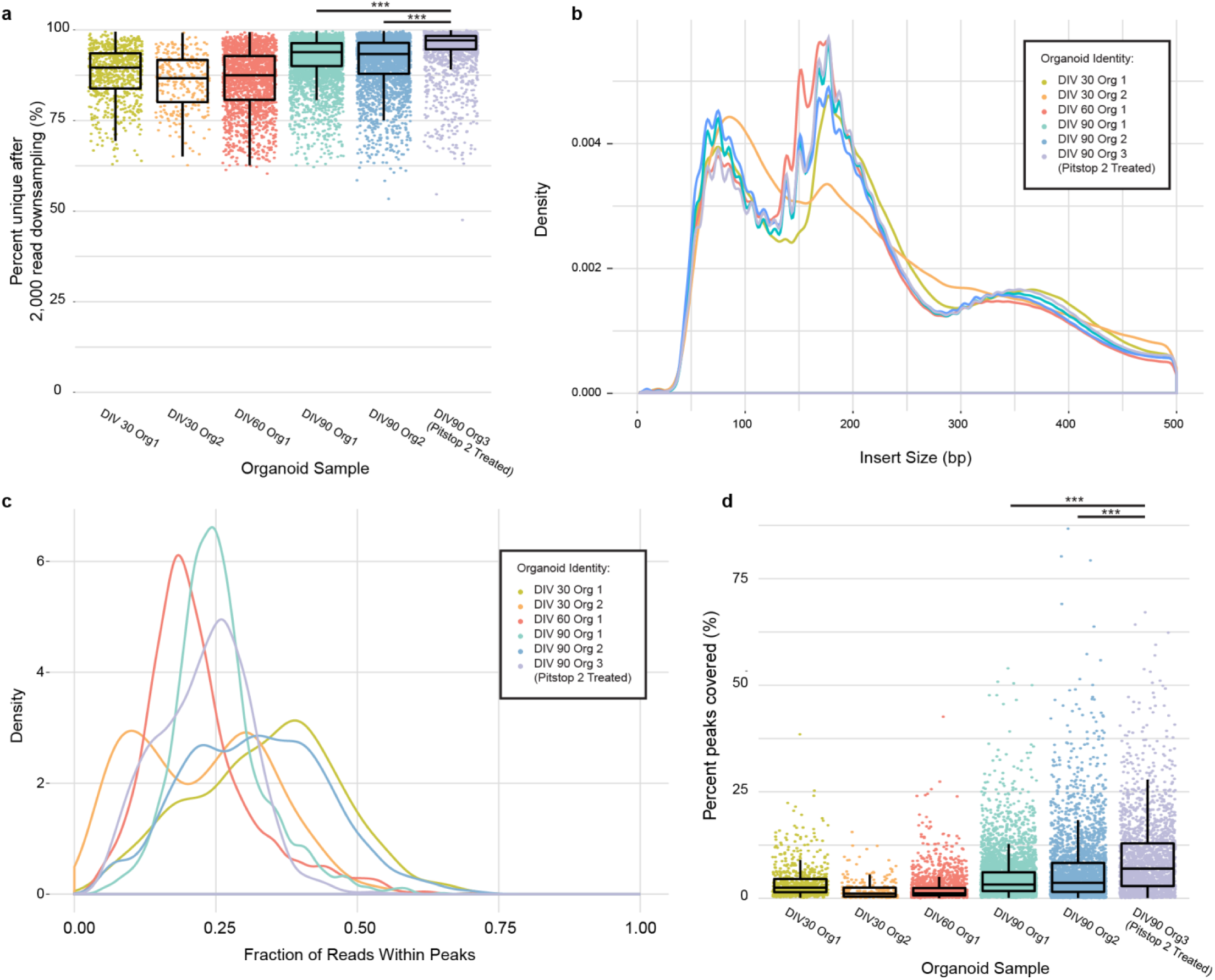
Single-cell library performance and complexity from sourced organoid samples. **a**, Bar plot of single-cell library complexities after down-sampling. Single-cell libraries were randomly sampled for 2,000 reads. The percent of unique reads per library is plotted. Significance was calculated by DIV90 Pitstop 2 70 μM treatment condition to the DIV90 0 μM Pitstop 2 treatment organoids with a two-sided Mann-Whitney test. **b**, Density plot of sequencing molecule insert sizes by treatment condition. **c**, Density plot of fraction of reads overlapping with *de novo* called peaks per condition. **d**, Estimated peak coverage per cell based on read count within *de novo* called peaks. Significance was calculated by DIV90 Pitstop 2 70 μM treatment condition to the DIV90 0 μM Pitstop 2 treatment organoids with a two-sided Mann-Whitney test.*** adjusted p-value ≤ 10^−4^

**Extended Data Fig 6.**
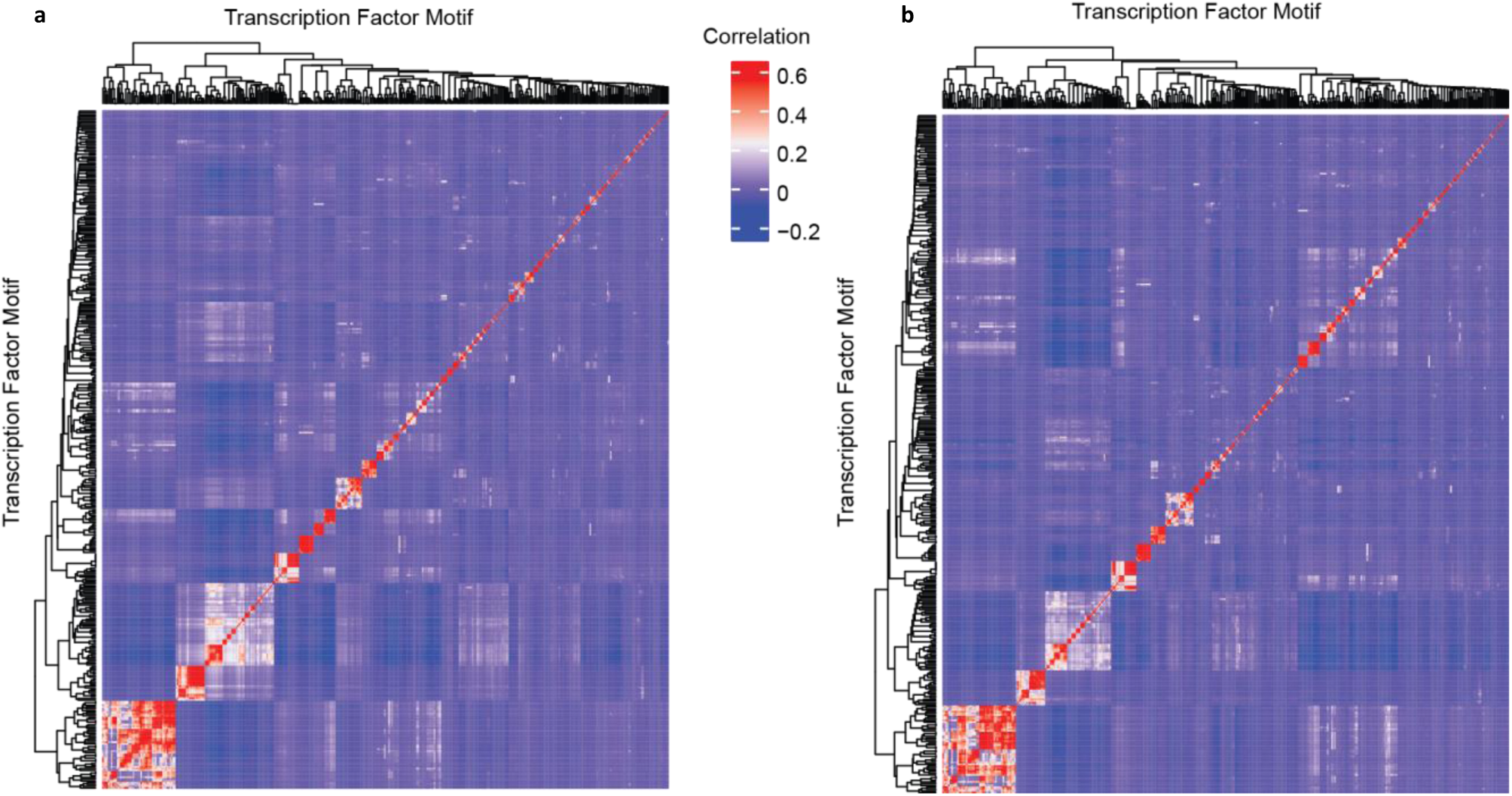
Transcription factor motifs share the same pattern of co-occurrence in novel and known peak regions. **a**, Hierarchically clustered heatmap of Pearson’s correlation between motif presence in known enhancer regions. **b**, Hierarchically clustered heatmap of Pearson’s correlation between motif presence in novel-defined enhancer regions.

**Extended Data Fig 7.**
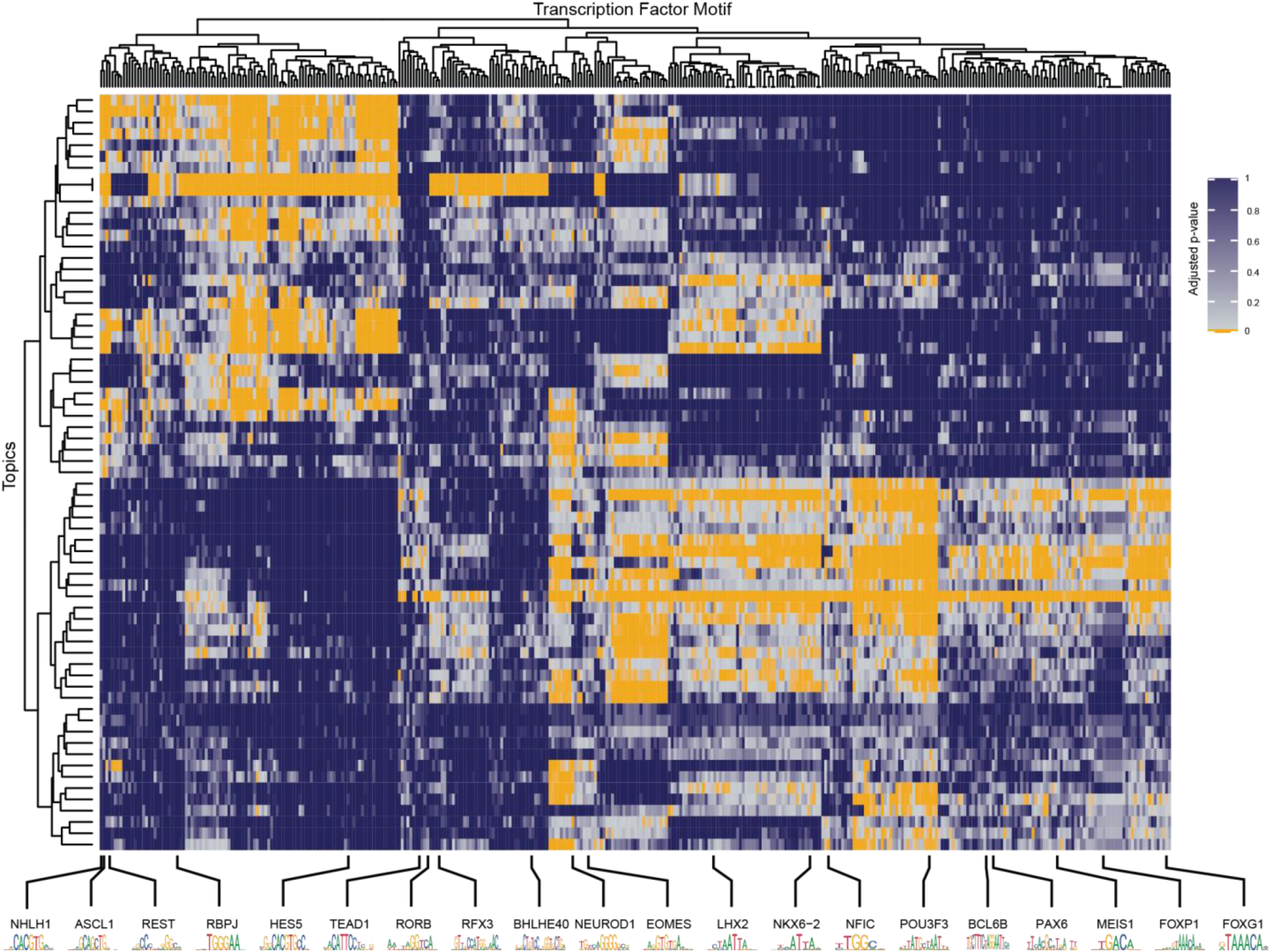
Topic Characterization. Heatmap of topic-assigned peak accessibility enrichment for transcription factor motifs. Values were calculated using *chromVAR* for motif assignment per peak, followed by hypergeometric test compared to the set of all peaks within topics. p-values were adjusted for multiple testing by Benjamini-Hochberg correction. Rows and columns of heatmap were hierarchically clustered by Pearson’s correlation. Select transcription factors and associated motifs are shown.

**Extended Data Fig 8.**
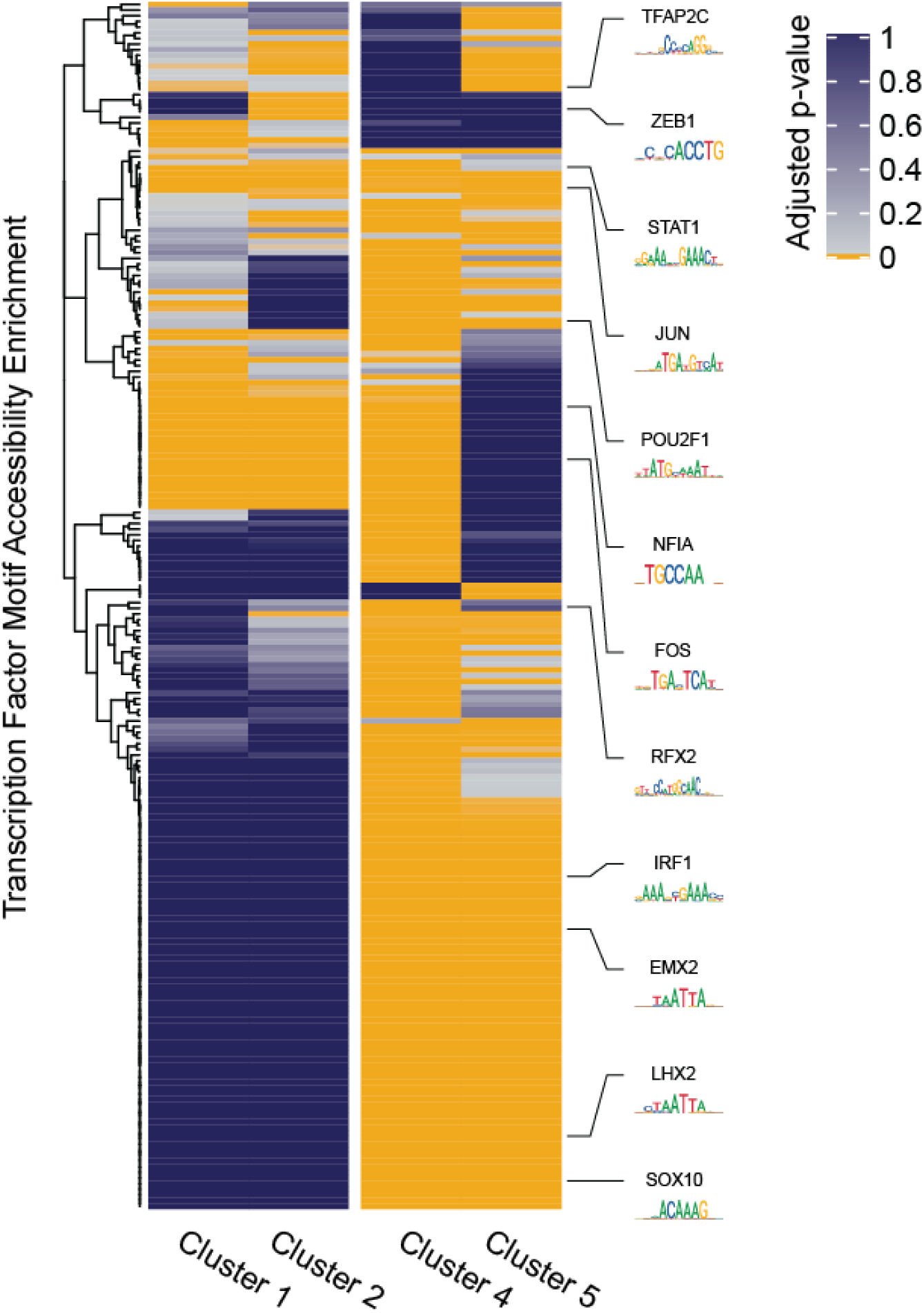
Cluster 3 shows transcription factor motif enrichment in outer radial glia markers in pairwise comparison to other clusters. Heatmap of topic-assigned peak accessibility enrichment for transcription factor motifs using differentially accessible peaks generated through cluster 3 to individual other clusters. Values were calculated using *chromVAR* for motif assignment per peak, followed by hypergeometric test compared to the set of all peaks within topics. P-values were adjusted for multiple testing by Benjamini-Hochberg correction. Rows of the heatmap were hierarchically clustered by Pearson’s correlation. Select transcription factors and associated motifs are shown.

**Extended Data Fig 9.**
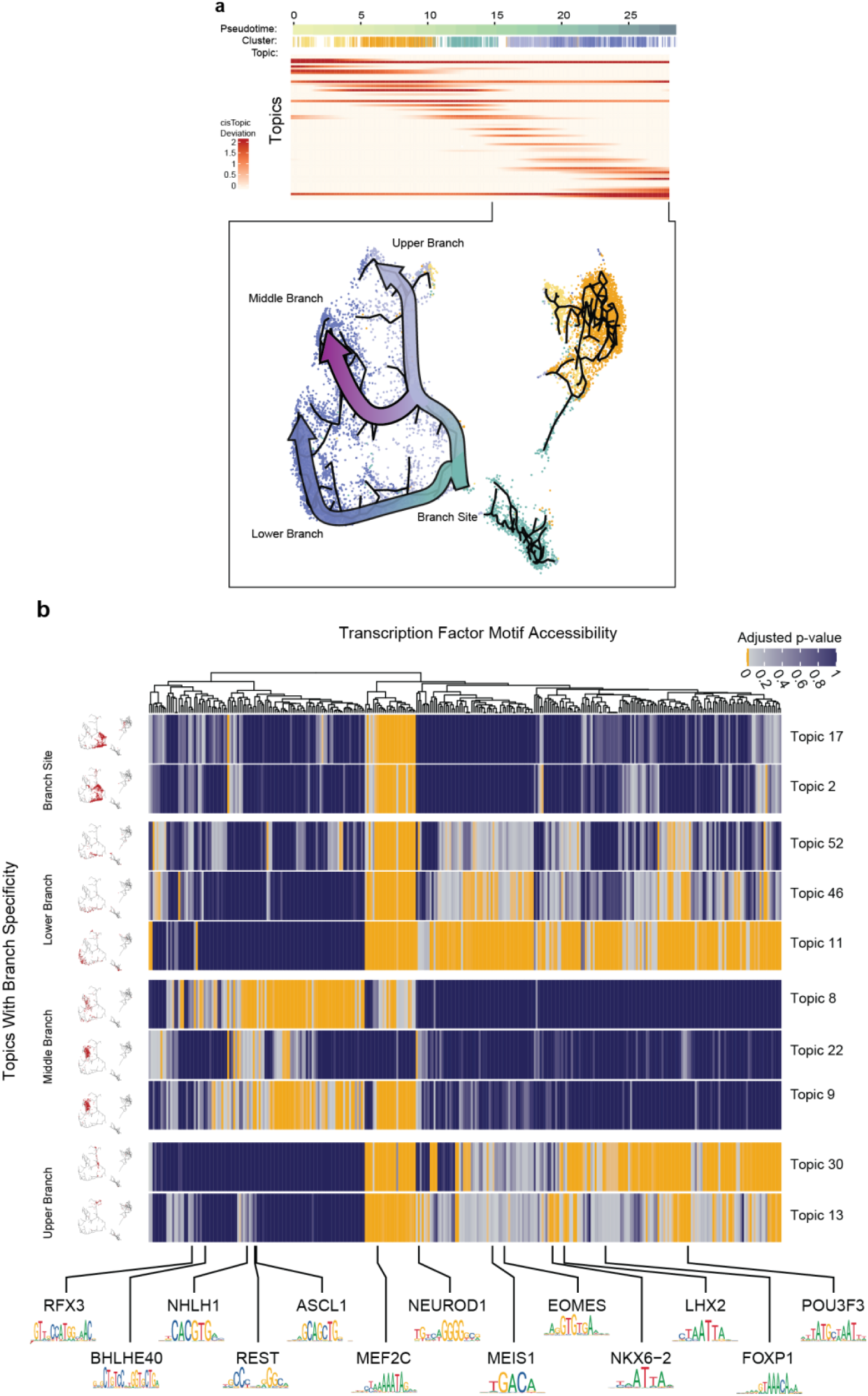
Topics through pseudotime. **a**, (Upper) Topic accessibility smoothed by LOESS fit and ordered through pseudotime by maximum value. (Lower) Schematic showing major branches within clusters 4 and 5 with maximum accessibility occurring at pseudotime appearance of cluster 4. Major branches are categorized by max accessibility split by upper/middle/lower branches and the branch site. Cluster assignment coloring is consistent with Fig. 2c. **b**, Subset of topics maximally accessible during the branch point of the neurogenic lineage. (Left) Cells colored by highest accessibility per topic (Z-score ≥2 colored red, other cells omitted) used to classify topic branch specificity. (Right) Transcription factor motif accessibility per topic. Topics ordered by branch grouping and pseudotime. Heatmap of topic-assigned peak accessibility enrichment for transcription factor motifs. Values were calculated using *chromVAR* for motif assignment per peak, followed by hypergeometric test compared to the set of all peaks within topics. P-value was adjusted for multiple testing by Benjamini-Hochberg correction. Select motifs and associated JASPAR motifs are shown below.

**Extended Data Fig 10.**
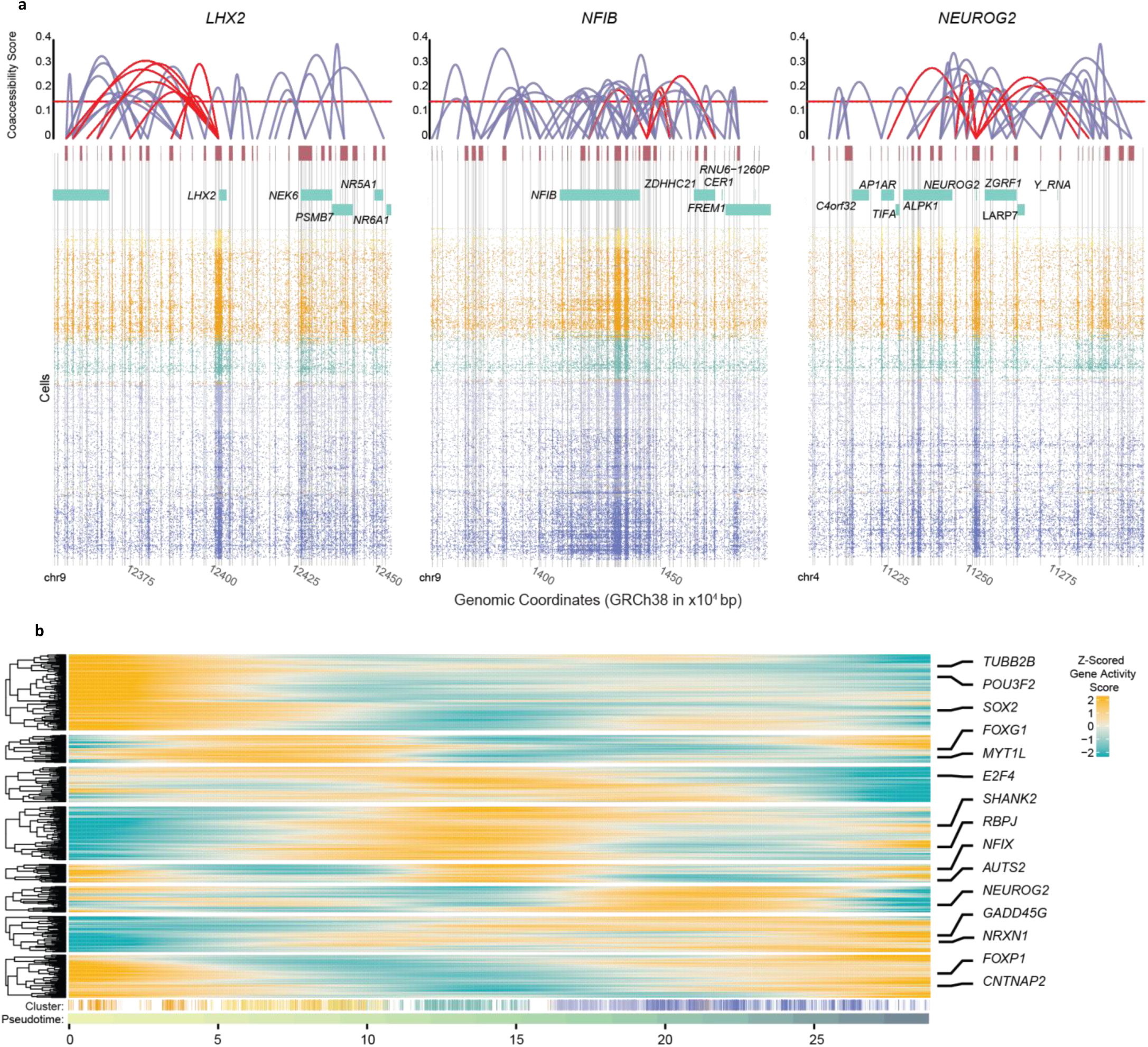
Cicero coaccessibility networks and gene activity scores through pseudotime. **a**, Example coaccessibility scores and peak linkages generated through cicero (top). A cut-off coaccessibility score of 0.15 was used for generation of gene activity (GA) scores (shown as a red horizontal bar). Cis-coaccessibility networks (CCANs) specific for the example genes are highlighted with red peak connections. Underneath each co accessibility plot is aligned aggregated peaks (red boxes), annotated genes (blue) and called peak regions (gray vertical bars). Read plots were aligned underneath for the same genomic region. **b**, LOESS smoothed plot of Z-scored gene activity (GA) ordered through pseudotime. Rows were hierarchically clustered and separated into 8 clusters for visualization. Selected genes are shown on the right. Cluster assignment coloring is consistent with Fig. 2c.

## Online Methods

### Sample generation

#### U2OS Cell culture

U2OS human osteosarcoma cells (ATCC, Cat. HTB-96) were maintained at 37°C and with 5% CO_2_ in culture medium comprised of DMEM (Thermo Fisher, Cat. 11995065) supplemented with 10% fetal bovine serum (FBS; VWR, Cat. 89510-186). For imaging, cells were plated in #1.5 cover glass bottom 8-well chamber slides (Thermo Fisher Scientific, Cat. 155409) at a density of 1×10^5^ cells per well and grown for 2 days prior to fixation and labeling for the ATAC-see protocol^14^.

#### GM12878 cell culture

The human B-lymphocyte line GM12878 was obtained from the NIGMS Human Genetic Cell Repository at the Coriell Institute for Medical Research. GM12878 cells were cultured in 5% CO _2_ at 37°C. Cells were grown in Roswell Park Memorial Institute media (RPMI, Thermo Fisher, Cat. 11875093) supplemented with 15% (v/v) FBS (Thermo Fisher, Cat. 10082147), 1x L-glutamine (Thermo Fisher, Cat. 25030081), 1x penicillin-streptomycin (Thermo Fisher, Cat. 15140122), and 1x gentamicin (Thermo Fisher, Cat. 15750060). Cells were grown to ∼70% confluency within a 25 mL flask at the time of harvest.

#### iPSC culture

Forebrain organoids were differentiated from a human control iPSC line (CW20043) obtained from the California Institute for Regenerative Medicine (CIRM) repository at the Coriell Institute for Medical Research. The iPSCs were maintained feeder-free on 100 mm^2^ dishes coated with 5 μg/mL vitronectin (Thermo Fisher, Cat. A14700) in StemFlex Medium (Thermo Fisher, Cat. A3349401) and kept inside a 37°C incubator with 4% O_2_ and 5% CO_2_. Prior to passaging or thawing, StemFlex Medium was supplemented with the following ROCK inhibitors to promote viability: 10 μM Y27632 (Stemgent; Cat. 04-0012-10) and 1 μM Thiazovivin (Stemgent, Cat. 04-0017). The culture medium was exchanged daily on weekdays; a double volume of media was provided on Fridays. The iPSCs were passaged at ∼80-90% confluency as follows: cells were washed once with PBS then incubated with Versene (Thermo Fisher, Cat. 15040066) for 4 minutes at 37°C. After Versene was aspirated, StemFlex medium containing 10 μM Y27632 and 1 μM Thiazovivin was added to the culture dish to collect cells. The iPSC suspension was further diluted in StemFlex with Y27632 and Thiazovivin then transferred to new vitronectin-coated dishes using a 1:12-1:25 split ratio depending on colony density prior to passaging. CryoStor10 freezing medium (STEMCELL Technologies, Cat. 07930) was used for cryopreservation of iPSCs.

#### Differentiation of forebrain organoids

Differentiation of forebrain organoids from human iPSCs was carried out using a previously described protocol with certain modifications^4,40^. In this modified protocol, cortical neurogenesis is initiated in AggreWell800 plates (STEMCELLTechnologies; Cat. 34850) to produce uniformly-sized embryoid bodies that are chemically induced to develop into forebrain-like tissues through the addition of small molecule SMAD inhibitors. Refer to Supplementary Table 2 for the regimen of factors to add to the differentiation medium used during feedings. AggreWell800 plates were prepared for aggregate culture according to manufacturer instructions using STEMdiff Neural Induction Media (NIM; STEMCELL Technologies, Cat. 05835) supplemented with 10 μM Y27632, 2 μM Thiazovivin, 2 μM Dorsomorphin (Tocris, Cat. 3090), and 2 μM A83-01 (Tocris, Cat. 2939) then set aside until needed. Of note, all supplemental factors (e.g. small molecules, recombinant proteins) added to differentiation medium were done so immediately prior to feeding.

To begin neural induction (day 0 of *in vitro* differentiation; DIV 0), two 80-90% confluent 100 mm^2^ dishes of iPSCs were washed once with 10 mL PBS then treated with 5 mL Accutase (STEMCELL Technologies, Cat. 07920) for 8 minutes at 37°C. Colonies were disaggregated into a single cell suspension by pipetting up and down for ∼10 seconds with a 5 mL serological pipet, then for an additional 3-5 seconds with a P1000 pipet. The single iPSCs in Accutase were transferred to 50 mL conical tube then the 100 mm^2^ dishes were immediately washed twice with 10 mL of DMEM/F12 with GlutaMax (Thermo Fisher, Cat. 10565018) that was added to the same 50 mL tube containing cells. Residual clumps of cells were removed by passing the suspension through a 40 μm cell strainer into a new 50 mL conical tube. Filtered iPSCs were centrifuged at 200 × g for 5 minutes, after which, the supernatant was aspirated and the pellets resuspended in 1 mL NIM containing 10 μM Y27632, 2 μM Thiazovivin, 2 μM Dorsomorphin, and 2 μM A83-01. Cells were kept on ice while counting and performing calculations.

Each well in the AggreWell800 plate contains 300 microwells. Each of these microwells is used to form a single organoid initially comprised of approximately 10,000 cells. To achieve this, we seeded 3,000,000 iPSCs per well of the AggreWell800 plate, centrifuged the plate at 100 × g for 3 minutes to collect ∼10,000 cells into each of the 300 microwells, then incubated at 37 °C with 5% CO_2_. Organoids were cultured in the AggreWell800 plate for 5 days with a daily 75% medium change. On day 2, Y27632 and Thiazovivin were omitted from the differentiation medium. On day 5, organoids were harvested from the AggreWell800 plate according to manufacturer instructions using wide-bore P1000 tips that were prepared by cutting off tips with a clean pair of scissors then autoclaved to sterilize. Organoids collected from a single well of the AggreWell800 plate were transferred to one 60 mm^2^ ultra-low attachment dish (Corning, Cat. 3261) in Forebrain Differentiation Medium I (FDM I) comprised of DMEM/F12 with GlutaMax, 1% N2 Supplement (Thermo Fisher, 17502048), 1% MEM Non-Essential Amino Acids (Thermo Fisher, Cat. 11140050), 1% Penicillin/Streptomycin (Thermo Fisher, Cat. 15140122), and 1 μg/mL heparin (Sigma, Cat. H3149) supplemented with 1 μM CHIR99021 (Stemgent, Cat. 04-0004) and 1 μM SB431542 (Stemgent, Cat. 04-0010). On day 6 (the following day), organoids were embedded in 20 μL droplets of Growth Factor Reduced (GFR) Matrigel (Corning, Cat. 354230) using wide-bore P200 tips as previously described^41^, then returned to ultra-low attachment suspension culture in FDM I with 1 μM CHIR99021 and 1 μM SB431542 and for another 8 days with medium changes every 2 days.

On day 14, GFR Matrigel-embedded organoids were transferred to culture in either a nylon (laser sintered) or ULTEM 9085m 3D-printed Spin Omega 12-well miniature bioreactor^4,40^ (refer to Qian *et al*^4^. for detailed instructions on bioreactor 3D-printing and assembly) in Forebrain Differentiation Medium II (FDM II) consisting of a 1:1 mix of DMEM/F12 with GlutaMax and Neurobasal (Thermo Fisher, Cat. 17504044) with 1% N2 Supplement, 2% B27 Supplement (Thermo Fisher, Cat. 17504044), 0.5% GlutaMax (Thermo Fisher, Cat. 35050061), 1% MEM Non-Essential Amino Acids, 1% Penicillin/Streptomycin, 2.5 μg/mL Insulin (Sigma, Cat. I9278), and 50 μM 2-Mercaptoethanol (Sigma, Cat. M3148) with medium changes every 2-3 days. Approximately every two weeks, organoids were transferred to different wells in a new 12-well bioreactor plate to avoid position effects^4,40^. On day 70, we began adding 0.2 mM L-Ascorbic Acid (Sigma, Cat. A4403), 0.5 mM cAMP (Sigma, Cat. A9501), 20 ng/mL BDNF (Peprotech, Cat. 450-02), and 20 ng/mL GDNF (Peprotech, Cat. 450-10) to the FDM II and continued exchanging the medium every 2-3 days up until day 90 when the experiment ended.

#### Organoid freezing protocol

Single organoids were transferred to individual 1.5 mL tubes using a P1000 pipet equipped with a wide-bore tip then pelleted by centrifugation at 500 × g for 2 minutes at 4 °C. After removing the supernatant, organoid pellets were flash frozen by placing the tubes in a slurry of ethanol and dry ice for approximately 2 minutes then transferred to a −80 °C freezer for storage.

### Pitstop 2 increased transposome nuclear occupancy

We reasoned that disruption of the nuclear pore complex (NPC) would allow for increased permeability to the ∼56 kDa transposomes used in ATAC-seq protocols. To test this, we used two orthogonal modifications to loaded oligonucleotides to assess changes in the early stages of the ATAC-seq protocol. The two methods were performed based off analysis done in the ATAC-see protocol and early steps of flow-assisted nuclei sorting (FANS) in the sci-protocols^6,9,14^.

#### Generation of fluorescent transposome complex for FANS

Transposase nuclear occupancy in a large population of cells was measured by use of Cy5-labelled transposome. We generated a Cy5-modified mosaic end oligonucleotide (Cy5-ME; /5Phos/CTGTCTCTTATACACATCT/3Cy5Sp/, IDT) and loaded naked transposase enzymes following previously described protocols^42^. i5 and i7 indexed tagmentation oligonucleotides (i5Transposase and i7Transposase, respectively) were annealed to the Cy5-ME in equimolar ratios at 100 μM in the 2X transposase dialysis/dilution buffer^42^. Annealing was performed on a thermocycler with the following conditions: 95°C for 5 min, then a stepwise decrease in temperature of 5°C every 2 minutes until reaching 20°C. 16 μM each of the annealed i5Transposase:Cy5-ME and i7Transposase:Cy5-ME oligonucleotide complexes were placed into a 96-well plate, resulting in a final concentration of 8 μM annealed transposome species per well. 8 μM transposase was added suspended in 500 nM NaCl (Fisher, Cat. M-11624) added to each well. The mixture was assembled by incubation at 25 °C for 1 hour and stored at −20°C until ready for use.

#### Nuclei isolation of GM12878

To isolate nuclei for the GM12878 line, 10 mL of suspended cells were taken from the 25mL flask (See “GM12878 cell culture” above). The 10 mL suspension was spun at 4°C at 500 rcf for 10 minutes to pellet. Media was aspirated and the pellet was resuspended in 500 μL ice cold PBS (VWR, Cat. PI283448) to wash. The cell suspension was then pelleted again by a 500 rcf centrifugation at 4°C for 5 minutes. Liquid was aspirated and the pellet resuspended in 1 mL of Nuclei Isolation Buffer (NIB; 10mM Tris HCl, pH 7.5 [Fisher, Cat. T1503 and Fisher, Cat. A144], 10mM NaCl [Fisher, Cat. M-11624], 3mM MgCl2 [Sigma, Cat. M8226], 0.1% IGEPAL [v/v; Sigma, I8896], 0.1% Tween-20 [v/v, Sigma, Cat. P7949] and 1x protease inhibitor [Roche, Cat. 11873580001]). Nuclei were incubated on ice for at least 5 minutes, and held on ice until further processing. Nuclei were quantified on a hemocytometer and diluted to 2,000 nuclei/μL with further addition of NIB.

#### Pitstop 2 nuclei treatment and flow assisted nuclei sorting (FANS)

To quantify the additive effect of Pitstop 2 (Abcam, Cat. ab120687) on transposomes nuclear occupancy, we tested several concentrations during nuclear isolation and tagmentation: 0 μM, 50 μM, 70 μM and 90 μM. The various conditions were treated the same throughout the protocol, with the exception of nuclear isolation and the tagmentation reaction buffers. To modify the nuclear isolation condition, 2X versions of NIB+Pitstop 2 (NIB with Pitstop2 at 0 μM, 100 μM, 140 μM or 180 μM concentrations) were added in equal volume to aliquots of the 2,000 nuclei/μL. This diluted samples to 1,000 nuclei/μL. All conditions were stored on ice throughout the addition of the modified NIB. To modify the tagmentation reaction, 2X TD buffer (Illumina, Cat. 15027866) aliquots were prepared with a supplementation of 3 mM Pitstop 2, to the appropriate final concentrations (0 μM, 50 μM, 70 μM and 90 μM). 50 μL of the appropriately supplemented 2X TD Tagment DNA buffer was added to 50 μL (50,000 nuclei total) samples, respective of test condition (100 μL final volume). For each reaction, 5μL of 8 μM loaded Cy5 fluorescent transposase (See “Generation of fluorescent transposome complex for FANS” above) was added. Tagmentation was carried out simultaneously for each reaction by incubation at 55°C for 15 minutes with gentle mixing (300 rpm) on thermomixer. Following this, all reactions were placed immediately on ice. Nuclei were pooled and subsequently spun down at 500 rcf for 10 minutes at 4°C. Tagmented nuclei were washed with 500 μL NIB and run through a 35 μm cell strainer (BD Biosciences, Cat. 352235). 3μL of DAPI (5mg/mL) was added to tagmented nuclei samples. Each sample was then run through a SONY SH800 cell sorter for quantification of DAPI and Cy5 fluorescence.

#### Preparation of fluorescent probes for microscopy

To mimic conditions of the previously published ATAC-See paper^14^, Cy3 labeled oligonucleotide probes with complementary sequences to the single-stranded overhangs of the NEX1A and NEX2A adaptor oligonucleotides (Illumina, Inc.) were synthesized by Integrated DNA Technologies (IDT). The oligonucleotide sequences are as follows: NEX1A_rev-Cy3, 5’-GACGCTGCCG/3Cy3Sp/-3’; NEX2A_rev-Cy3, 5’-CCGAGCCCACG/3Cy3Sp/-3’.

#### ATAC-see labeling of nuclei and Imaging

The protocol for ATAC-see labeling of nuclei was adapted from Chen et al^14^. Cells cultured in 8-well chambers (See “U2O2 cell culture”) were briefly washed with 1× PBS (VWR, Cat. PI283448) before light fixation with 1% formaldehyde (FA) at room temperature (RT) for 10 min, followed by washing with 1× PBS for 5 min. The cells were then permeabilized with either Nuclei Isolation Buffer (NIB; 10mM Tris HCl, pH 7.5 [Fisher, Cat. T1503 and Fisher, Cat. A144], 10mM NaCl [Fisher, Cat. M-11624], 3mM MgCl2 [Sigma, Cat. M8226], 0.1% IGEPAL [v/v; Sigma, I8896], 0.1% Tween-20 [v/v, Sigma, Cat. P7949] and 1× protease inhibitor [Roche, Cat. 11873580001]) or NIB with 70 μM Pitstop 2 (final concentration) at RT for 10 min and rinsed with 1× PBS (2×5 min). Next, the samples were blocked in 5% bovine serum albumin (BSA) (Sigma-Aldrich, Cat. A7906) at RT for 15 min and rinsed with 1× PBS. A transposase mix containing 60 μL of 2× Tagment DNA (TD) buffer and 2 μL transposase adapter complex (TDE1) were added to each well; both the TD buffer and the TDE1 complex were from the Nextera DNA Sample Preparation Kit (Illumina Inc., Cat. 15027865, 15027866). For our Pitstop 2 treated nuclei condition, TD buffer was supplemented with 70 μM Pitstop2 (final concentration).The samples were incubated at 55°C for 15 min with gentle mixing (300 rpm) on a thermomixer for the transposition reaction to complete. After the reaction, the samples were washed with 1× PBS containing 0.01% SDS and 50 mM EDTA at 55 °C (3× 15 min), cooled to room temperature, and fixed again with 3.7% fresh paraformaldehyde (PFA, v/v, Sigma, 158127) for 10 min. The samples were then stained in 1.5 μM DAPI (ThermoFisher Scientific, Cat. D1306), washed with 1× PBS (3×5min) and stored in 1× PBS at 4 ^°^C until imaging.

The TDE1 transoposome complexes were nonfluorescent. To visualize the transposome complexes that have reacted with genomic DNA, the two fluorescent oligonucleotide probes, namely NEX1A_rev-Cy3 and NEX2A_rev-Cy3, were added to TDE1-labeled samples after dilution in the hybridization buffer, Buffer C^43^ (500 mM NaCl in 1× PBS) to a final concentration of 25 nM. The mixture was incubated at RT for 30 min to allow for efficient hybridization between the Cy3-oligonucleotide probes to their respective adapters (NEX1A and NEX2A), which contain single-stranded overhangs with complementary sequences to the probes, on the transposome complex (TDE1). This step yields specific labeling of the TDE1 complexes. Residual Cy3-oligonucleotide probes were removed by aspiration. The samples were then rinsed with and left in Buffer C for imaging on a Zeiss Laser Scanning Microscope (LSM 880) located at the Advanced Light Microscopy Core (ALMC) in Oregon Health and Science University (OHSU).

Imaging with the LSM 880 was conducted using a 63x oil immersion objective (Plan-Apochromat 63x/1.4 Oil DIC) using the 405 nm and 561 nm lasers for DAPI and Cy3 excitation, respectively, with emission bands for each channel set to default values for the two fluorophores in the Zeiss software.

### scip-ATAC-seq for Pitstop 2 concentration optimization

#### GM12878 nuclei isolation, Pitstop 2 treatment and tagmentation

To quantify the additive effect of Pitstop 2 (Abcam, Cat. ab120687) on library complexity, we tested several concentrations: 0 μM, 50 μM, 70 μM and 90 μM. Nuclei isolation for GM12878 cells and NIB+Pitstop 2 supplementation were done as described above. All conditions were stored on ice throughout the addition of the modified NIB. To modify the tagmentation reaction, 2X TD buffer (Illumina, Cat. FC-121-1031) aliquots were prepared with a supplementation of 3 mM Pitstop 2, to the appropriate final concentrations (0 μM, 50 μM, 70 μM and 90 μM). 5μL of the appropriately supplemented 2X TD buffer was added to 5 μL (5,000 nuclei total) samples, respective of test condition (10 μL final volume). We prepared 192 tagmentation reactions (two 96-well plates). GM12878 cells were split with the following numbers per condition: 32 reactions with 0 μM Pitstop 2, 64 50 μM Pitstop 2 reactions, 64 70 μM Pitstop 2 reactions and 64 90 μM Pitstop 2 reactions. For each reaction, 1μL of 8 μM loaded indexed transposase was added (See Picelli *et al.* for loading protocol)^42^. Tagmentation was carried out as described above. All reactions were pooled respective of Pitstop 2 conditoin and 3μL of DAPI (5mg/mL) was added to each.

#### Sorting nuclei

One 96-well plate was prepared, each well containing 8.5 μL of protease buffer (PB; 30 mM Tris HCl, pH 7.5 [Fisher, Cat. T1503 and Fisher, Cat. A144], 2 mM EDTA [Ambion, Cat. AM9261, 20 mM KCl [Fisher, Cat. P217 and Fisher, Cat. A144], 0.2% Triton X-100 [v/v, Sigma, Cat. 9002-93-1], 500 ug/mL serine protease [Fisher, Cat. NC9221823). To each well, a combination of 1 μL 10 mM i5 and 1μL 10 mM i7 PCR primers (Supplementary Table 8, IDT) containing a well-specific index combination was added. DAPI-stained nuclei pools were then sorted using a Sony SH800 FACS machine with sample and sorting chambers held at 5°C. Gating was performed to isolate a clean population of singlet nuclei. Each well within the two prepared 96-well plates had 44 nuclei deposited.

#### Transposase denaturation and PCR

Following sorting, plates were spun at 500 rcf for 5 minutes at 4°C to ensure nuclei were within the reaction buffer. Transposomes and any other proteins within the reaction were then degraded by the serine protease by holding the samples at 55°C for 20 minutes, the protease was then denatured by heating to 70°C for 30 minutes. Following this, 13.5 μL of PCR Master Mix (13 μL 2X KAPA Hotstart HiFi [Fisher, Cat. NC0465187], 0.25 μL (2 units) Bst3.0 [NEB, Cat. M4374] and 0.25 μL 100X SYBR Green I [FMC BioProducts, Cat. 50513]) was added to each well. Real-time (RT)-PCR was performed on a BioRad CFX machine, for the following temperatures and times: 72°C for 5 minutes, 98°C for 30 seconds, and then multiple rounds of 98°C for 30 seconds, 63°C for 30 seconds, 72°C for 1 minute and a SYBR plate read before starting the next cycle. PCR reactions were stopped when the SYBR readout for a majority of wells plateaus (18-22 cycles).

#### Library pooling and cleanup

For the PCR plate, 10 μL of each well was pooled for clean-up and quantification. First, the 960 μL pool was concentrated on a PCR purification column (Qiagen, Cat. 28106) following manufacturer’s protocol. DNA was eluted off the column in 50 μL 10mM Tris HCl, pH 8.0 (Fisher, Cat. T1503 and Fisher, Cat. A144). Library pools were then cleaned and size selected using SPRI beads generated as describe previously^9^. An equal volume of prepared SPRI beads (1X) were added to the library pools, and incubated at room temperature for 15 minutes. Beads were then pelleted on a magnetic rack and washed twice with 150 μL of freshly prepared 80% ethanol (v/v, Decon, Cat. 2705). Following the second wash, all remaining liquid was carefully removed from the tube without disrupting the pellet. Pellets were then allowed to dry for 10 minutes, before being resuspended in 50 μL 10mM Tris HCl, pH 8.0. This 1:1 volume SPRI bead clean-up was repeated a second time. For the final elution of the second cleanup, libraries were eluted in 27 μL 10mM Tris HCl, pH 8.0. 2 μL of this eluate was used for quantification with a Qubit HS Assay (Thermo Fisher, Cat. Q32851). Following this, libraries were diluted to 4 ng/μL and 1μL was run on an Agilent DNA HS BioAnalyzer (Agilent, Cat. 5067-4626). Libraries were then diluted based on BioAnalyzer-reported molarity in the 150-1000bp range and loaded on a NextSeq 500 (Illumina Inc.) sequencer High Capacity kit with a loading concentration of 1.2 pM with a custom sequencing protocol, for 75 cycles of read 1, 30 cycles of index 1 and index 2, and 75 cycles of read 2^13^.

### Organoid characterization

#### Immunohistochemistry

Methods used to prepare organoids for cryosectioning were adapted from a previously described protocol^41^. In brief, wide-bore P200 or P1000 tips were used to transfer two to three organoids to single wells in a 24-well plate containing 250 μL medium for each cryosection block to be embedded. Organoids in the 24-well plate were washed once with 1 mL PBS then fixed with 1 mL of 4% PFA (Sigma, Cat. 158127) for 15 minutes at 4°C. Organoids were then washed three times with 1 mL PBS for 10 minutes at room temperature. The final PBS wash replaced with 30% sucrose (Sigma, Cat. S7903) with 0.02% sodium azide (Sigma, Cat. S2002) then the plate was incubated for 24-72 hours at 4°C. Subsequently, organoids were equilibrated in 1 mL of a 7.5% gelatin (Sigma; Cat. G1890)/10% sucrose embedding solution inside a 37°C incubator for 15-30 minutes. During this period, biopsy cryomolds (Tissue-Tek, Cat. 4565) for each well with organoids by coating the bottoms with a thin layer of the gelatin/sucrose embedding solution. Organoids were transferred to the cryomolds pre-coated with embedding solution using wide-bore tips then incubated at 4°C for 5 minutes. Cryomolds were then filled with embedding solution and allowed to solidify for 20-30 minutes at 4°C. The gelatin/sucrose blocks with organoids were frozen in a −30°C to −50°C isopentane bath for 2 minutes, after which, the cryomolds with frozen blocks were tightly wrapped with Parafilm and stored at −80°C until sectioned.

For immunohistochemistry, frozen blocks with embedded organoids were cut into 20 μm sections using a cryostat (Leica). Sections were serially collected onto Superfrost Plus microscope slides (Fisherbrand, Cat. 22-037-246), allowed to dry for ≥30 minutes, and then stored at −20°C or 4°C until ready for staining. Prior to staining, slides were equilibrated to room temperature then sections were circumscribed with a hydrophobic barrier pen (Invignome, Cat. GPF-VPSA-V). Sections were washed twice with PBS for 10 minutes then blocked for 1 hour at room temperature in permeabilization/blocking buffer comprised of PBS with 10% normal goat serum (Jackson ImmunoResearch, Cat. 005-000-121), 1% bovine serum albumin (BSA, Millipore, Cat. 126626), 0.3% Triton X-100 (TX-100, Sigma, Cat. 11332481001), 0.05% Tween-20 (Sigma, Cat. P1379), 0.3 M glycine (Sigma, Cat. G7126) and 0.01% sodium azide (Sigma, Cat. S2002) for 1 hour at room temperature. During the blocking step, primary antibodies (Supplementary Table 3) were diluted in a buffer containing PBS, 2% NGS, 1% BSA, 0.01% TX-100, 0.05% Tween-20, and 0.01% sodium azide. The diluted primary antibodies were applied to sections then incubated overnight at 4°C inside a StainTray (Simport Scientific, M922-2). Primary antibodies were washed from the sections five times with PBS for 5 minutes at room temperature. During wash steps, secondary antibodies (Supplementary Table 4) were prepared by diluting 1:1000 in the same buffer used to dilute primary antibodies. Sections were incubated with the diluted secondary antibodies inside a StainTray for 1 hour in the dark at room temperature (sections were protected from light following secondary staining). Secondary antibodies were washed from the sections three times with PBS for 5 minutes, then nuclei were counterstained with DAPI (Thermo Fisher, Cat. D1306) for 10 minutes at room temperature. After DAPI staining, sections were washed an additional two times then glass coverslips were mounted with ProLong Diamond Anti-Fade Mounting Medium (Thermo Fisher, Cat. P36961).

#### Microscopy and image processing

Live organoid images (Extended Data Fig. 3) were taken with a Nikon Ts2 inverted microscope. Optical sections of organoids (Fig. 2a; Extended Data Figs. 3 and 4) were acquired with a Zeiss ApoTome AxioImager M2 fluorescent upright microscope and processed using Fiji software.

#### Organoid sci-ATAC-seq and scip-ATAC-seq

The sci-ATAC protocol for organoids was similar to that performed for testing Pitstop 2 concentrations with the following exceptions. 1) Nuclei were isolated from the flash frozen pellets by resuspension with NIB. For DIV 30 organoids, 300 μL NIB was used; for DIV 60 and DIV 90 organoids, 600 μL of NIB was used. Resuspension was done by 10-20 triturations of NIB solution to break up the pellet of cells. Cells were then incubated on ice for 10 minutes and then triturated another 10 times. The full volume of each sample was then each run through a 35 μm cell strainer (BD Biosciences, Cat. 352235) and nuclei were stained with 3 μL of DAPI (5mg/mL, Thermo Fisher, Cat. D1306) for DIV30 samples, and 5 μL DAPI for DIV60 and DIV90 samples. To demonstrate library complexity improvement, one DIV 90 organoid sample was dissociated with 70 μM Pitstop2 NIB solution, prepared as described above. 2) A Sony SH800 FACS machine was used to sort 5,000 nuclei (identified through DAPI gating) into each well of multiple 96 well plates. The wells containing sorted nuclei (n=432 wells) were then tagmented in parallel as described above (Supplementary Table 8). The nuclei treated with Pitstop 2 prior to sorting were once again treated with a final concentration of 70 μM Pitstop 2 added to the TD buffer. 3) 100 tagmented nuclei were deposited into each well of the PCR plates by FANS sorting due to the higher number of tagmentation reactions and thus lower chance of barcode collisions. 4) All wells were PCR amplified for 19 cycles. Libraries were cleaned and quantified as described above and sequenced on the Illumina NextSeq 500, and HiSeq™ 4000.

### Computational Analysis

#### FACS quantification of transposase nuclear occupancy

Flow cytometry data was analyzed by exporting fcs formatted files from the Sony SH800 sorter. These files were then processed by use of the flowCore package in R (v1.48.1, v3.5.1, respectively)^44^. All conditions were first gated to include singlet nuclei through DAPI staining. Following this, Cy5-Height fluorescence was measured for nuclei singlets (n = 32,511, 34,008, 15,500, 16,440 and 12,890 nuclei for non-fluorescent transposase control, 0 μM Pitstop 2, 50 μM Pitstop 2, 70 μM Pitstop 2 and 90 μM Pitstop 2 treatment conditoins, respectively) and plotted by the ggridges package (v0.5.1) in R.

#### Fluorescent transposome imaging analysis

Image analysis was performed using custom Matlab (Mathworks, MA) scripts using the image processing toolbox. First, cell nuclei were identified and segmented by applying Gaussian filters to raw, grayscale DAPI images followed by a Sobel-Feldman operator^45^ for nuclei edge detection. Binary masks for whole nuclei were created with image opening operation and verified visually. Around 40-50 nuclei were randomly chosen from all acquired images from both Pitstop 2 treated and non-treated populations for further analysis.

To quantify how signal intensity varies with respect to the distance from the nuclear periphery, a serial erosion was performed on each nuclei using the binary masks. Each erosion shrunk the mask in all directions by 4 pixels, and the difference image between the current and the last mask yields a 4-pixel wide annulus with a distance of (n-1)*4 + 2 pixels to the nuclear periphery in all directions, where n is the current iteration of erosion. The series of annular marks were applied to the original grayscale image to calculate the mean pixel intensity across each annulus, which was then normalized to that of the outer most three annuli. The normalized pixel intensities were plotted with respect to the corresponding distances to the nuclear periphery. Two-sided Mann-Whitney U tests and Benjamini-Hochberg p-value adjustment was performed in R via the *wilcox.test* function for each erosion in order to determine the statistical significance of the distribution of Cy3 normalized signal between Pitstop 2 treated and non-treated nuclei.

3D rendering of z-stack images was conducted with the Imaris software (Bitplane) using shadow projection and other relevant features. The interior features of the nuclei were visualized using the clipping plane tool (Fig. 1 b,c; Extended Data Fig. 1).

#### FastQ generation, index assignment, single-cell read set definition

Following sequencing libraries on both Illumina NextSeq 500 and HiSeq™ 4000 machines, bcl files were converted to FastQ format using bcl2fastq (v2.19.0, Illumina Inc.) with the option “with-failed-reads”. FastQ reads were then combined across sequencing runs. Read sets were allocated to index barcodes with an allowance of two Hamming distance from possible index combination (perl v5.16.3, custom script). FastQ-format reads were modified so as to have the accepted indexing barcode (cell ID) as the read name. FastQ files were then aligned to the hg38 reference genome (GRCh38, NCBI) via bwa-mem (v0.7.15-r1140)^46^. The resulting aligned reads then underwent a removal of duplicate reads based on unique cell ID, chromosome and start sites of reads. The number of total reads and persisting unique reads was used for a comparison of library complexity (unique reads/total reads respective of cell ID). Single-cell libraries were defined by cell ID read sets containing a unique read cut-off of at least 2,000 reads with Q>=10 mapping quality. In total we generated 12,076 sci(p)-ATAC-seq libraries. This yielded a single-cell library generation efficiency of 12,076/43,200 or 27.95% of the theoretical maximum for our experiment’s sci-ATAC-seq/scip-ATAC-seq strategy and sorting schema.

#### Pitstop 2 library complexity assessment

To quantify the effects of Pitstop 2 treatment on sci-ATAC-seq/scip-ATAC-seq library complexity, several analyses were performed on the GM12878 samples. Reads were allotted to cell ID read pools by demultiplexing and aligned to the human reference genome (hg38) as described above. A Mann-Whitney U test was used as a non-parametric statistic for library complexity comparison and Benjamini-Hochberg correction for the p-value is reported. To account of uneven sequencing effort, cell IDs which passed cutoff (and were thus determined to be single-cell libraries) were subsampled randomly to 2,000 total reads. The percentage of unique reads per cell was used to generate a measure of population library complexity per Pitstop 2 treatment condition. Again a Mann-Whitney U test with Benjamini-Hochberg correction was used. Both full cell ID complexity and subsampled cell ID complexities were plotted using the *geom_jitter* and *geom_boxplot* function dependent on Pitstop 2 treatment condition by the *ggplot* package (Fig. 1, Extended Data Fig. 2; v3.1.0). The read length distribution was generated by use of the uniquely aligned paired-end sequencing data and plotted using the *geom_density* function in ggplot. The fraction of reads on target characterization was generated in the following way: Following the definition of putative single-cell libraries, we used libraries passing filter to generate a pseudo-bulk sample. We performed *de novo* peak calling on this pseudo-bulk sample using macs2 (v2.1.1.20160309) with default settings^47^. This was to assess putative regions of open chromatin by read pileup^5^. Peak sizes were expanded to 500 bp windows, with the center being designated as narrowPeak site defined by macs2. The percentage of uniquely aligned reads which overlapped with peak regions was used to calculate the fraction on target (FRIP) value for each cell. The distribution of FRIP values was plotted dependent on condition using *geom_density* and a Mann-Whitney U test with Benjamini-Hochberg correction was used to assess differences (Extended Data Fig. 2; Supplementary Table 1). The same methods for quality control metrics were used for analysis of Organoids sci-ATAC-seq library complexities respective of each cell’s organoid origin (Fig. 2, Extended Data Fig. 5; Supplementary Table 5).

#### Peak calling and characterization

For scip-ATAC-seq/sci-ATAC-seq organoid samples, we called peaks as read-pileup regions (described above). In total, we uncovered 125,645 peaks. To characterize peaks in the context of existing data sets, we used the *chromVAR* R package to find transcription factor motif occurrence in each peak, using the JASPAR set of motifs (v1.4.1, v2018 release, respectively)^31,39,48^. We also looked for overlapping regions within the H3K27ac ChIP-seq defined enhancer regions within human cortex, or organoid samples defined by the PsychENCODE Consortium^16,21^ and other enhancer regions collated by the Genehancer data base^20^ and overlapped them with our peak set through the use of bedtools (Supplementary Table 6, v2.22.0)^49^. We compared known enhancer regions (those which overlapped either PsychENCODE or Genehancer data sets) to novel peaks by comparing the co-occurrence of motifs within peaks. We used the *chromVAR* characterization of all JASPAR motifs within all peaks, subset by novel or known enhancer region. We then took each pairwise transcription factor binding motif comparison (for example, the occurrence of the NEUROD1 DNA binding motif and TBR1 binding motif both within the same peak), and used a Pearson’s correlation to assess how likely the two motifs are to occupy the same peak. We then generated a Pearson’s correlation between the two resultant square matrices to assess motif-motif cooccurence in peaks globally. This informed the bias of called peaks for DNA motifs, and suggested that novel peaks had similar motif binding site characteristics to known enhancer regions. We clustered motif co-occurrence respective of novel/known region using the *Heatmap* function in *ComplexHeatmap* with hierarchical clustering of the Pearson’s correlation for visualization (Extended Data Fig. 6, v1.20.0;)^50^.

#### Generation of counts matrix and cisTopic dimensionality reduction

We used the filtered 125,645 peaks from our data set and the cell ID-associated, deduplicated reads to generate a cell ID by peak read count matrix, wherein each element within the matrix describes the number of reads from the respective cell ID overlapping with the peak feature. For each single-cell library an average of 26.59 ± 10.74 % (Extended Data Fig. 5c, mean ± s.d.) of reads overlapped with peaks. Cell IDs were filtered to have a minimum of 500 non-zero peak elements in the matrix. Peak features were then filtered to have a minimum of 25 cell IDs with non-zero values. This filtered to a final count of 10,355 cells by 125,645 peak features matrix. We then performed feature selection through the *cisTopic* R package (v0.2.0)^22^. Briefly, single-cell ATAC-seq counts matrices are a highly sparse, with read-dropout and chromatin inaccessibility limiting information. The cisTopic algorithm measures the correlation between peaks to define broader “topics” which suffer less noise than individual peaks. We performed cisTopic analysis with the default settings on the binarized, filtered cell ID by peak features matrix. We determined that a count of 67 topics was optimal for our data structure, based on maximum log-likelihood values. Each topic had maximally accessible peaks extracted by use of the cisTopic *binarizecisTopics* function with a p-value threshold of 0.975, yielding a median of 2762 peaks per topic.

To characterize topics, we used their respective peak region motif annotations attained through *chromVAR* as described above for peak characterization. We performed a hypergeometric test for each motif and each topic, and performed a Benjamini-Hochberg correction. Motifs with no topic peak sets falling below an adjusted p-value of 0.001 were discarded. The remaining motifs were plotted by *Heatmap* function (ComplexHeatmap, v.1.20.0, R)^50^. We observed transcription factor motif enrichment in peak sets suggesting they may be capturing biological information (Extended Data Fig. 7).

#### Cell clustering

We performed Louvain clustering via the R package *Rphenograph* on the full cisTopic matrix with the following options: n-neighbors = 2000 (v0.99.1)^23^. This approach defined five clusters with membership numbers ranging from 922 to 2986 cells. For visualization and dimensionalty reduction, we performed uniform manifold approximation and projection (umap) via the R package *umap* on the full cisTopic matrix to cluster cells in three-dimensions (v0.2.0.0)^24^. We plotted the projection (first two dimensions) with various coloring schemes matching annotations for phonograph cluster assignment, organoid source, organoid source DIV, and Pitstop 2 treatment (Fig. 2c-f, respectively). We found an unequal proportion of DIV-sourced cells through the clusters, suggesting shifting cell type populations through differentiation (Fig. 2e, Supplementary Table 5).

#### Cell type assignment

Since the ATAC-seq signal is an indirect measurement of genomic active regions, we sought to define cell types within our clusters by differential accessibility (DA) of peaks between our five clusters. To define a DA peak set for each cluster, we first bolstered information content per cell by aggregating local groupings. We aggregated cells from our 3 dimensional projection, using k-means algorithm (base R) respective of cluster. We generated 657 cell aggregates (mean 15.6 cells per aggregate) for our 5 clusters (mean of 130.8 aggregates per cluster). These aggregates became our samples per cluster. We then performed differential analysis on each cluster by all other clusters (1 by 4) using the DESeq2 R package (v.2-1.22.2) on our aggregates using the Wald Test with a false discovery rate (fdr) correction. Peaks with a Log2 fold change ≥ 0.5 and an adjusted p-value ≤ 0.1 were defined as DA. 2,625, 67,691, 168, 13,102, and 6,120 peaks were defined as DA for cluster 1 through 5, respectively.

DA peaks were used to test for motif enrichment over the set of all DA peaks across clusters. To do this, we used the annotated the peak regions from *chromVAR*, described above for peak characterization. We performed a hypergeometric test for each motif and each cluster, and performed a Benjamini-Hochberg correction. Motifs with no cluster DA peak sets falling below an adjusted p-value of 0.001 were discarded. The remaining motifs were plotted by *Heatmap* function (Fig. 2g; ComplexHeatmap, v.1.20.0, R)^50^. Due to the lack of DA peaks statistically enriched in cluster 3, we lacked power for proper motif enrichment. We also performed cluster 1,2,4, and 5 by cluster 3 pairwise DA by the same method as the cluster by all other cluster comparison. 13,885, 1,996, 3,273, and 1,772 peaks were defined as DA between cluster 3 and clusters 1, 2, 4, and 5, respectively. Motif enrichment and plotting was performed in the same way (Extended Data Fig. 8).

Cell type markers used are defined by the cell type marker sets described in Nowakowski *et al.*^19^, filtered to those present within the JASPAR data base (2018 release, full list in Supplementary Table 7)^39^. Cell types were defined by three assumptions. First we looked at the motif enrichment in DA peaks for each cluster. We reasoned that active transcription factor families are more likely to have presented DNA binding motifs. It is notable that within a transcription factor family, many proteins may share the same predicted motif^39^. We noted enrichment of the NEUROD, T-box and MEF2 family of transcription factors in clusters 4 and 5. We observed enrichment of proliferative markers such as the Sox family or ESRRB, as well as forebrain specification linked markers in clusters 1 and 2 (PAX6, OTX2, ETV5, NKX6-2). Cluster 3 had few DA peaks, suggesting it was a middle state between clusters 1 and 2, and clusters 4 and 5. Second, we looked at loci-specific read density for corticogenic marker genes (Fig. 2g). We used the read density within the region, and the proportion of reads per cluster aligning with our called peaks as suggestive of gene activity. We generated read plots flanking the genic regions of SOX2, PAX6, EOMES, TBR1, and NEUROD2. Wherein each row of the plot represents a single cell, and each point represents a uniquely aligned read to that genomic region. We under-laid this plot with our called peaks (represented as gray vertical bars). We colored cell-read points to match the cluster coloring scheme within the same figure (Fig. 2h). And thirdly, we further accepted DIV-sourced cell proportion of each cluster as informative, wherein later born cell types in corticogenesis are suspected to have a higher proportion of later DIV sampled cells (Fig. 2e). Together, we conservatively defined clusters 1 and 2 as neuroepithelial to proliferating radial glia cells, cluster 3 as outer radial glia cells, and cluster 4 and 5 as intermediate progenitors to post-mitotic neurons.

#### Pseudotime and transcription factor activity through pseudotime

The cortical organoid cells are all generated from the same starting state, that is to say, the primed neuroepithelial-like induced pluripotent stem cells^4^. Since there is a common starting population, we sought to characterize the progression of chromatin states from the stereotyped cell type changes from neural progenitor to neuron. To do this, we used monocle (Monocle 3 alpha version, http://cole-trapnell-lab.github.io/monocle-release/monocle3/)^51^. We supplied our pre-generated umap dimensions within monocle for consistency. We generated a minimal spanning tree via the *SimplePPT* function in three dimensions traversing all five clusters, and set the root node to be a branch within the previously identified stem-like cluster 1 (Fig. 3a). With each cell ID assigned a pseudotime value based on location along the minimal spanning tree, we sought to compare chromatin changes to known biology. To describe transcription factor motif enrichment on a cell level, we used the *chromVAR* deviation scores.^31^ We generated deviation scores via the *computeDeviations* function for each cell for each JASPAR 2018 motif^39^. We Z-scored the deviation values via the base R *scale* function, and performed a LOESS smoothing of the chromVAR-reported deviation score by cell ID ordered by pseudotime by the *loess* function (100 bins, *loess* function from base R). We plotted the resulting smooth curves via *ComplexHeatmap* package^50^, clustering by Spearman’s correlation on rows (Fig. 3b).

We asked if the accessibility of topic peak sets also changes through pseudotime. We performed the same LOESS smoothing on the ordered array of cell topic values through pseudotime (Extended Data Fig. 9a). We found that many topics were pseudotime specific, and thus we selected a time point in which we observed branching along the minimal spanning tree to better characterize epigenetic changes. We categorized topics by cells with maximal (Z-score ≥ 2 topic deviation value) topic accessibility into branches along the minimal spanning tree coinciding with cluster 4 and 5. We then grouped these topics using the same transcription factor enrichment scores previously generated. We observed clear biases in branches, suggesting a dynamic epigenome (Extended Data Fig. 9b). To confirm DNA motif enrichment within topics by branch, we also plotted *chromVAR* reported motif accessibility with a binary cutoff of deviation-score ≥ 2.

#### Identification of coaccessible peaks using *Cicero*

Chromatin is a dynamic and tightly regulated structure. The use of nearby enhancer regions has an improved positive predictive value for transcription over promoter accessibility alone^35^. To leverage nearby accessibility peaks in this fashion, we employed the *cicero* package from the Trapnell group using default settings (v1.0.15)^35^. We generated cis-coaccessiblity networks using these coaccessbility scores (cut-off of coaccessibility scores ≥ 0.15; Fig. 3c; Extended Data Fig. 10a). We used these cis-coasccesbility scores to generate gene activity scores, a cumulative measure of promoter and enhancer regions anchored by peaks overlapping promoter regions. These gene activity scores were generated on aggregated cells (k=300 cells per aggregate, using k-nearest neighbors given the umap coordinates). This generated 1,235 aggregates. We used the mean pseudotime value of all 300 cells per aggregate as the aggregate pseudotime score. We then performed the same LOESS smoothing of gene activity scores through pseudotime on the aggregates (Fig. 3d, Extended Data Fig. 10b).

